# An Adhesive Interface for the Non-Clustered δ1 Protocadherin-1 Involved in Respiratory Diseases

**DOI:** 10.1101/498196

**Authors:** Debadrita Modak, Marcos Sotomayor

## Abstract

Cadherins form a large family of calcium-dependent adhesive proteins involved in morphogenesis, cell differentiation, and neuronal connectivity. Non-clustered δ1 protocadherins form a cadherin subgroup of proteins with seven extracellular cadherin (EC) repeats and cytoplasmic domains distinct from those of classical cadherins. The non-clustered δ1 protocadherins mediate homophilic adhesion and have been implicated in various diseases including asthma, autism, and cancer. Here we present X-ray crystal structures of Protocadherin-1 (PCDH1), a δ1-protocadherin member essential for New World hantavirus infection that is typically expressed in the brain, airway epithelium, skin keratinocytes, and lungs. The structures suggest a binding mode that involves antiparallel overlap of repeats EC1 to EC4. Mutagenesis combined with binding assays and biochemical experiments validated this mode of adhesion. Overall, these results reveal the molecular mechanism underlying adhesiveness of PCDH1 and δ1-protocadherins, also shedding light on PCDH1’s role in maintaining airway epithelial integrity, the loss of which causes respiratory diseases.

---

Cells are the basic units of life and selectively organize themselves to form tissues with the help of various adhesion proteins that engage in homophilic (same type) and heterophilic (different type) contacts. Cell-cell adhesion is of prime importance for the formation and maintenance of multicellular structures. Some of the main families of adhesion proteins include integrins, selectins, cadherins and members of the immunoglobulin superfamily^1–4^.

Cadherins are calcium-dependent adhesion glycoproteins involved in a variety of biological processes such as cell differentiation and tissue morphogenesis^5^, cell signaling^6–8^, mechanotransduction^9–11^, and brain morphogenesis and wiring^12–16^. These proteins have a unique structure including tandem extracellular cadherin (EC) repeats^5,17,18^, followed by a transmembrane and a cytoplasmic domain. The highly similar EC repeats are around 100 amino acids long and feature highly conserved calcium-binding motifs that bind three calcium ions at the linker regions between the repeats.

Non-clustered protocadherins are non-classical cadherins expressed in various tissues all over the body and have a wide range of functions. They are typically placed in two groups: δ and ε protocadherins^19^. The δ-protocadherins have seven (δ1) or six (δ2) EC repeats, a single transmembrane helix, and a cytoplasmic domain containing highly conserved motifs of unclear function (CM1, CM2 and CM3 in δ1; CM1 and CM2 in δ2)^19,20^. These play a major role in neuronal tissue connectivity and brain wiring and are involved in various neurological disorders such as epilepsy, autism and schizophrenia^20–22^.

Protocadherin-1 (PCDH1) is a non-clustered δ1 protocadherin involved in respiratory diseases. It was first identified by Sano and coworkers as PC42 in rat brain and retina tissue^23^. Later on, it was found to be mainly expressed in the airway epithelium, bronchial epithelium, lungs, skin keratinocytes and also in kidney, neural, and glial cells^24–28^. PCDH1 has seven EC repeats (labeled EC1 to EC7 from N to C-terminus), a single pass transmembrane domain and a cytoplasmic domain with the three conserved motifs (CM1-CM3) typical of the δ1 subgroup. Immunofluorescence microscopy experiments showed that L cells transfected with PCDH1 DNA aggregated in a calcium-dependent manner and expressed the protein along the boundary of the cells at cellular contact sites^23^. These results suggested that PCDH1 mediated aggregation through calcium-dependent homophilic interactions.

PCDH1 has been associated with asthma, a chronic inflammatory disorder of the airways characterized by wheezing, coughing, breathlessness, chest-tightness, bronchial hyperresponsiveness (BHR) and obstruction to airflow. It is a widespread health problem affecting people of all ages, especially children, and is caused by interaction of genetic and environmental factors^29,30^. The epithelial layer of the airway mucosa and bronchia acts as a barrier and prevents the passage of allergens and pathogens. Dysfunction of this epithelial layer barrier is thought to play a role in asthma^31^. PCDH1 may bind bronchial epithelial cells and epithelial cells of the airway mucosa together by mediating homophilic adhesion and helping in forming the epithelial barrier. Loss of adhesion by PCDH1 or underexpression of the protein may lead to loss of epithelial integrity and pathogenesis^31^. However, how PCDH1 and other members of the δ1-protocadherin family can mediate cellular adhesion is unclear.

PCDH1 has also been found to be essential for New World hantaviruses infection^32^. These viruses are transmitted from rodents to humans and cause a fatal respiratory disease called hantavirus pulmonary syndrome (HPS)^33,34^. There are no known vaccines or specific treatments for this disease in humans^35^. Results from *in vitro* and *in vivo* experiments indicate that direct recognition of PCDH1 EC1 by Gn/Gc glycoproteins on the virus envelope is a necessary step for infection^32^. However, the specific molecular mechanism by which PCDH1 EC1 interacts with the viral glycoproteins, and whether this interaction disrupts PCDH1 homophilic adhesion, remains to be elucidated.

The molecular mechanisms underlying adhesion by classical cadherins are well established. These proteins use their extracellular domains to form an intercellular (*trans*) bond that is mostly homophilic (between identical cadherins)^36,37^. The *trans*-homophilic bond is formed by contacts between the most membrane-distal EC repeats (EC1-EC1) coming from adjacent cells^38,36^. The EC1-EC1 contacts are mediated by the exchange of an N-terminal β-strand containing one or two tryptophan residues (Trp2 or Tpr2 and Trp4) that are docked into the hydrophobic pocket of the binding partner, thus forming a “strand-swapped” dimer.

Non-classical cadherins lack the tryptophans involved in classical adhesion and use different binding mechanisms. Cadherin-23 (CDH23) and protocadherin-15 (PCDH15) form a heterophilic bond essential for inner ear mechanotransduction^9,11^. The heterophilic CDH23-PCDH15 bond involves an antiparallel “extended handshake” arrangement that uses both EC1 and EC2 repeats^39^. In contrast, the clustered protocadherins (α, β, γ) and the non-clustered δ2 member PCDH19, all featuring six EC repeats, mediate *trans*-homophilic interactions using their first four extracellular repeats (EC1-4) arranged in an antiparallel bond, where EC1 interacts with EC4, EC2 with EC3, EC3 with EC2 and EC4 with EC1^40–45^. Missense mutations in PCDH19 that impair the formation of this bond are known to cause a form of epilepsy^42^.

The calcium-dependent adhesion mechanisms for various other non-classical cadherins, including the non-clustered δ1 protocadherins, are still poorly understood. Here we use bead aggregation assays to show that repeats EC1 to EC4 form the minimal adhesive unit for human (*Homo sapiens* [*hs*]) PCDH1. In addition, we present crystal structures of *hs* PCDH1 EC1-4 and show that an antiparallel overlapped dimer mediates *hs* PCDH1 adhesion. The discovery of this mode of adhesion, likely used by all other members of the δ1 protocadherin family, along with our structures, might help in understanding the molecular mechanisms underlying PCDH1-related respiratory diseases such as asthma and HPS.

## RESULTS

### Bead aggregation assays determine EC repeats required for adhesion

Previous studies have shown that the full-length PCDH1 protein mediates adhesion in cell-based assays^23^. To determine which EC repeats are essential for PCDH1 homophilic adhesion we created a library of constructs from the human protein (hereafter referred to as PCDH1) by deleting EC repeats from the C and N-terminus. Fc-tagged protein constructs were produced in HEK293T cells and binding assays were done with Protein G magnetic beads (see Methods). Results from bead aggregation assays of PCDH1 EC1-7Fc showed that PCDH1’s full-length extracellular domain mediates Ca^2+^-dependent homophilic adhesion (Fig. 1a). Aggregation was abolished in the absence of Ca^2+^ (2 mM EDTA) (Supplementary Fig. 1a). Next, to identify the minimum unit of homophilic adhesion mediated by the extracellular domain of PCDH1, bead aggregation assays were performed with PCDH1 EC1-6Fc, EC1-5Fc, EC1-4Fc, EC1-3Fc, EC1-2Fc, and EC2-7Fc (Fig. 1b-i). PCDH1 EC1-2Fc and EC1-3Fc showed no aggregation while PCDH1 EC1-4Fc, EC1-5Fc, EC1-6Fc, and EC1-7Fc showed Ca^2+^-dependent aggregation (Fig. 1 and Supplementary Fig. 1a-h). There was no aggregation when EC1 was truncated from the N-terminus (PCDH1 EC2-7Fc) (Fig. 1g). These experiments suggest that PCDH1 EC1-4 is the minimum unit of homophilic adhesion and that EC1 is required for adhesion.

**Figure 1.**
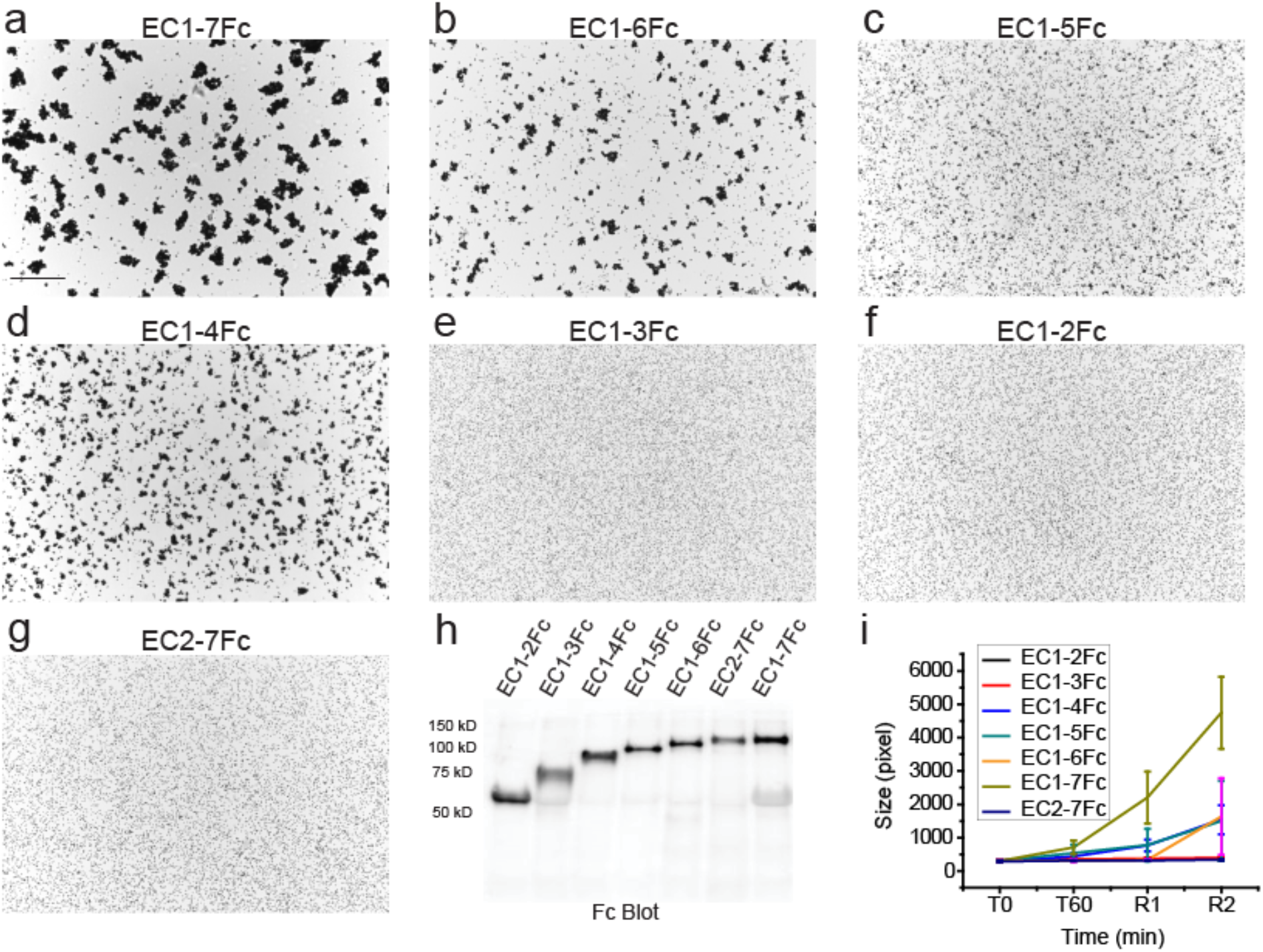
Minimal adhesive unit of PCDH1 is EC1-4. **(a-g)** Protein G beads coated with full length **(a)** and truncated versions **(b-g)** of the PCDH1 extracellular domain imaged after incubation for 1 hr followed by rocking for 2 min in the presence of calcium. Bar – 500 μm. **(h)** Western blot shows efficient expression of full length and truncated PCDH1 EC repeats. **(i)** Mean aggregate size for full length and truncated fragments of PCDH1 at T0 (*t* = 0 min), after 1 hr of incubation, T60 (*t* = 60 min) followed by rocking for 1 min (R1) and 2 min (R2). Error bars are standard error of the mean (*n* = 4 for all aggregation assays and constructs except for PCDH1 EC1-7Fc; *n* = 3).

### Bead aggregation and binding assays show that a disulfide-bond independent dimer mediates adhesion

In parallel to bead aggregation assays carried out with protein from HEK293T cells, we used PCDH1 EC1-4 and PCDH1 EC1-5 fragments refolded from *E. coli* inclusion bodies for binding assays that required large sample amounts. Refolding resulted in two monodisperse peaks corresponding to a monomer and a dimer mediated by an intermolecular cysteine-disulfide bond (data not shown). To test if free cysteines were necessary for PCDH1 dimerization, cysteine residues (C375 and C548) that were presumed to not be involved in intramolecular disulfide bonds were mutated to serine, thus abolishing potential PCDH1 intermolecular disulfide bonds. Bead aggregation assays performed with PCDH1 EC1-7Fc C375S C548S showed that PCDH1 mediated bead aggregation independent of these cysteines (Fig. 2a-b). Therefore, to avoid confounding effects all binding assays of PCDH1 EC1-4 and EC1-5 obtained from bacteria were done with proteins carrying the C375S (EC1-4, EC1-5) and C548S (EC1-5) mutations.

**Figure 2.**
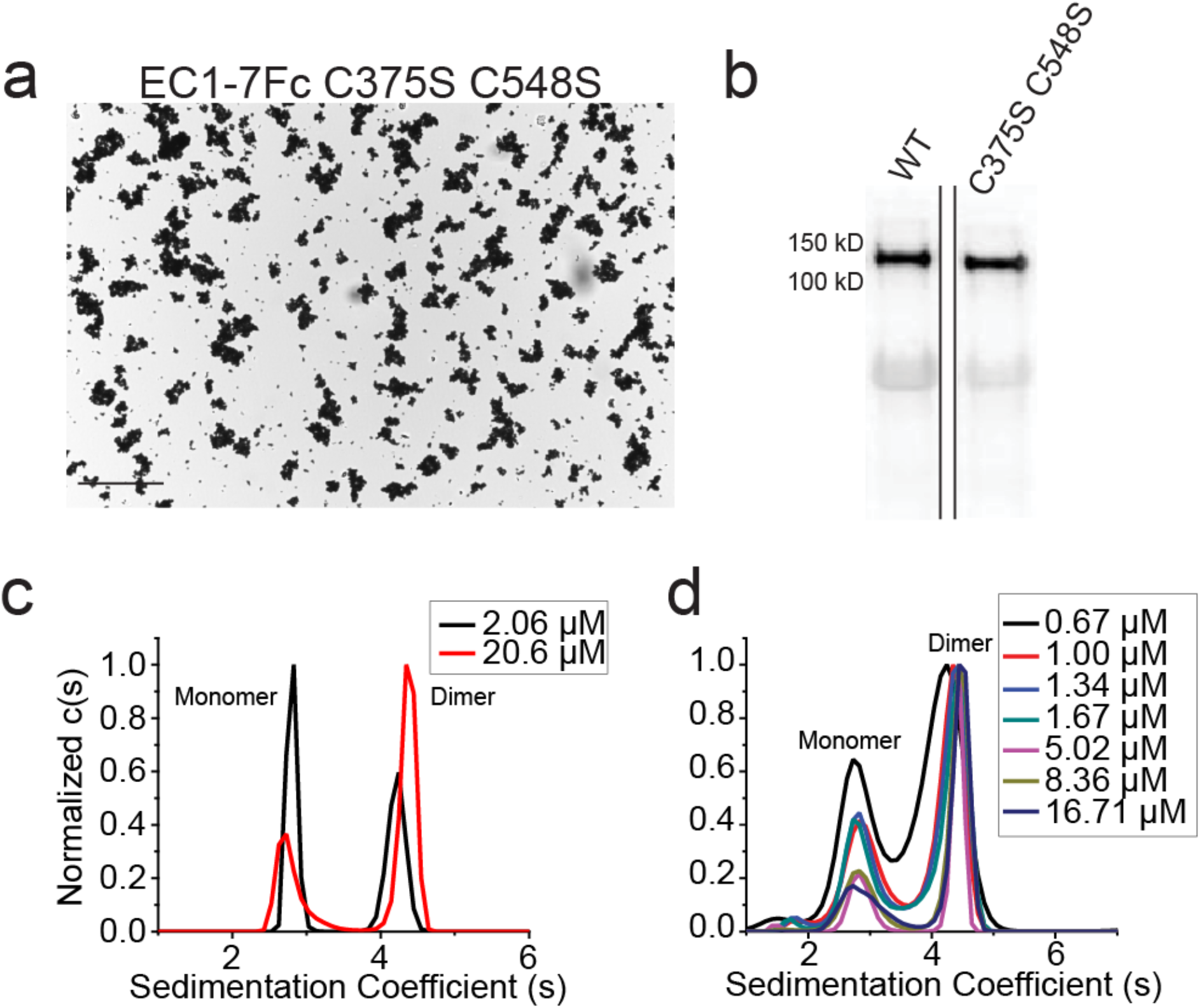
PCDH1-mediated bead aggregation is independent of disulfide bond formation and PCDH1 EC1-4 C375S and PCDH1 EC1-5 C375S C548S form dimers in solution. **(a)** Protein G beads coated with the PCDH1 EC1-7Fc C375S C548S mutant imaged after incubation for 1 hr followed by rocking for 2 min in the presence of calcium. Bar – 500 μm. Bead aggregation is evident. **(b)** Western blot shows efficient expression of wild type PCDH1 EC1-7Fc and mutant PCDH1 EC1-7Fc C375S C548S. **(c)** AUC data from bacterially produced PCDH1 EC1-4 C375S show both dimeric and monomeric peaks at 2.06 μM and 20.6 μM with normalized c(s) plotted against sedimentation coefficient (s). **(d)** AUC data from bacterially produced PCDH1 EC1-5 C375S C548S show both dimeric and monomeric peaks for a range of concentrations between 0.67 μM and 16.71 μM.

Analytical ultracentrifugation (AUC) sedimentation velocity experiments with a range of concentrations of PCDH1 EC1-4 C375S showed both monomeric and dimeric states in solution. The ratio of monomer to dimer was dependent on the concentration (Fig. 2c). Concentration dependent monomeric and dimeric states were also observed in sedimentation velocity experiments of PCDH1 EC1-5 C375S C548S (Fig. 2d). Samples were collected after AUC experiments and were run on SDS-PAGE, which indicated lack of significant degradation of the protein fragments (Supplementary Fig. 2a-b). Dimerization was observed for both PCDH1 EC1-4 C375S and PCDH1 EC1-5 C375S C548S fragments with the dimer dissociation constant (*K*_D_) estimated to be less than 5 μM. These results show that a dimer not formed by disulfide bonds mediates bead aggregation and exist in solution.

### X-ray crystal structures reveal overall architecture of PCDH1

Human PCDH1 protein fragments produced by bacterial and mammalian cells were used for crystallization and structure determination attempts. Two crystal structures were determined through molecular replacement and fully refined. The first one, PCDH1 EC1-4bc at 2.85 Å, was obtained using bacterially produced protein. The second, PCDH1 EC1-4mc at 3.15 Å, was obtained using protein produced by mammalian cells (Fig. 3 and Table 1). Both structures had one molecule in the asymmetric unit with good-quality electron density maps that allowed for the unambiguous positioning of side chains for most of the residues (Supplementary Fig. 3a-b). These included V3 to N443 in PCDH1 EC1-4bc and V3 to V439 in PCDH1 EC1-4mc, with both missing 8 residues in their EC1 repeat, and with PCDH1 EC1-4bc missing 4 residues in its EC4 repeat. The root mean square deviation (RMSD) for the Cα atoms among the chains of the two structures was ~1 Å. Since protomers were highly similar, we will describe features as seen in the PCDH1 EC1-4bc structure, unless otherwise explicitly stated.

**Table 1.** Statistics for *hs* PCDH1 structures.

| **Data collection** | ***Hs* PCDH1 EC1-4bc** | ***Hs* PCDH1 EC1-4mc** | ***Hs* PCDH1 EC3-4** |
| --- | --- | --- | --- |
| Space group | P6<sub>1</sub>22 | P6<sub>1</sub>22 | P4<sub>2</sub>2<sub>1</sub>2 |
| Unit cell parameters |  |  |  |
| *a*, *b*, *c* (Å) | 146.98, 146.98, 147.69 | 147.23, 147.23, 149.37 | 125.57, 125.57, 45.29 |
| $\alpha$, $\beta$, $\gamma$ (°) | 90, 90, 120 | 90, 90, 120 | 90, 90, 90 |
| Molecules per asymmetric unit | 1 | 1 | 1 |
| Beam source | 24-ID-E | 24-ID-E | 24-ID-E |
| Date of data collection | 18-DEC-2016 | 15-JUN-2017 | 30-JUL-2016 |
| Wavelength (Å) | 0.97921 | 0.97918 | 0.97921 |
| Resolution (Å) | 2.85 | 3.15 | 3.05 |
| Unique reflections | 22567 | 17231 | 7363 |
| Completeness (%) | 100.0 (100.0) | 100.0 (99.9) | 99.3 (90.6) |
| Redundancy | 10.7 (10.6) | 36.4 (28.2) | 9.9 (6.5) |
| *I* / $\sigma$(*I*) | 18.3 (1.923) | 58 (2.5) | 12 (1.67) |
| *R*<sub>merge</sub> | 0.156 (1.627) | 0.078 (1.762) | 0.166 (1.027) |
| *R*<sub>meas</sub> | 0.164 (1.710) | 0.079 (1.792) | 0.175 (1.110) |
| *R*<sub>pim</sub> | 0.050 (0.521) | 0.013 (0.321) | 0.054 (0.401) |
| *CC*<sub>1/2</sub> | 0.998 (0.716) | 1.004 (0.794) | 0.938 (0.551) |
| *CC* * | 0.999 (0.913) | 1.001 (0.941) | 0.981 (0.843) |
| **Refinement** |  |  |  |
| Resolution range (Å) | 48.26-2.85 (2.93-2.85) | 48.53-3.15 (3.22-3.15) |  |
| *R*<sub>work</sub> (%) | 22.1 (33.8) | 21.5 (33.3) |  |
| *R*<sub>free</sub> (%) | 24.3 (37.2) | 26.3 (43.0) |  |
| Protein Residues | 429 | 429 |  |
| Ligands/ions | 12 | 9 |  |
| Water molecules | 25 | 0 |  |
| Rms deviations |  |  |  |
| Bond lengths (Å) | 0.010 | 0.0095 |  |
| Bond angles (°) | 1.484 | 1.503 |  |
| *B*-factor average |  |  |  |
| Protein | 78.75 | 152.67 |  |
| Ligand/ion | 87.18 | 127.24 |  |
| Water | 47.34 | 0 |  |
| **Ramachandran Plot Region (PROCHECK)** |  |  |  |
| Most favored (%) | 87.8 | 83.2 |  |
| Additionally allowed (%) | 11.7 | 16.8 |  |
| Generously allowed (%) | 0.5 | 0.0 |  |
| Disallowed (%) | 0.0 | 0.0 |  |
| **PDB ID code** | **6BX7** | **6MGA** |  |

**Figure 3.**
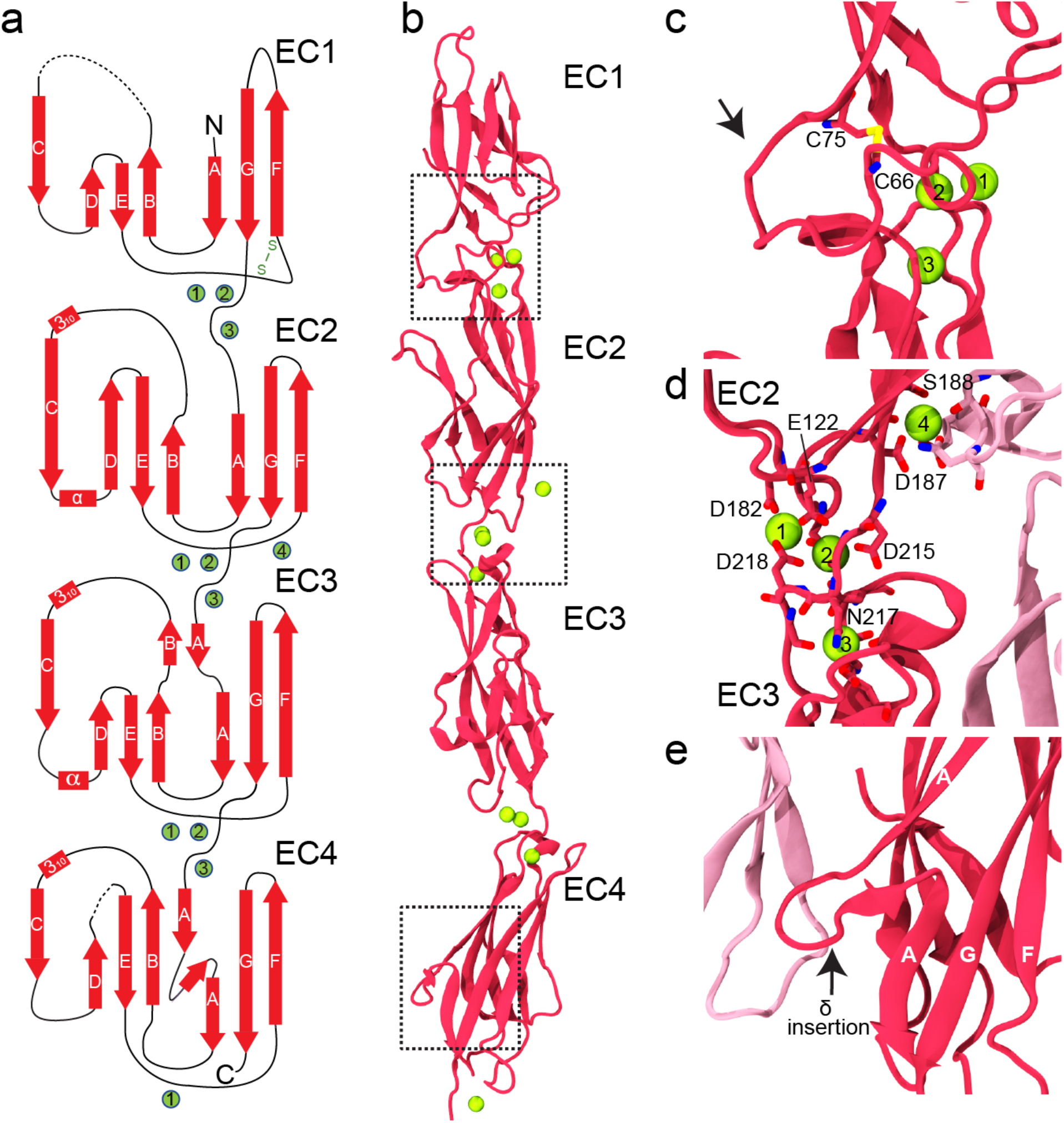
X-ray crystal structure of PCDH1 EC1-4. **(a)** Topology diagram of PCDH1 EC1-4bc. A typical cadherin fold is observed for each EC repeat with seven β strands labeled A to G. Calcium ions are shown as green circles. Dashed lines are missing loops. **(b)** Ribbon diagram of PCDH1 EC1-4bc (6BX7). Black dashed boxes indicate unique features of PCDH1 EC1-4 highlighted in panels (c-e). **(c)** Details of a disulfide loop in EC1 (arrow). **(d)** Details of the calcium-binding site in the EC2-EC3 linker region. **(e)** Details of an extended δ insertion within β strand A in EC4 not seen in clustered protocadherins (arrow).

PCDH1 EC1-4 has four tandemly arranged EC repeats with similar architecture (seven β strands labeled A to G arranged in a Greek-key motif; Fig. 3a-b). Although PCDH1 EC1-4 is somewhat straight as compared to the classical cadherins, which are curved, the EC repeats have the same overall classical architecture. There are three calcium ions in each complete linker region (EC1-EC2, EC2-EC3 and EC3-EC4) and one calcium ion at the end of EC4. Calcium ions at complete linker regions are coordinated by highly conserved canonical calcium-binding motifs XEX^BASE^, DXD, DRE/DYE, XDX^TOP^ and DXNDN (Supplementary Fig. 4). An additional calcium ion near the EC2-EC3 linker is unique and is at a crystal contact, but might be present due to purification and crystallization conditions with high calcium (Fig. 3b,d). The PCDH1 EC1-4mc structure does not have this additional calcium ion.

Another notable feature observed in the PCDH1 EC1-4 structure is the presence of a 10-residue long (C66-C75) disulfide loop in EC1, between β strands E and F (Fig. 3c). Shorter disulfide loops (7 residues) have been previously seen in the clustered and δ2 protocadherin structures at this location. The presence of an extended insertion with eleven residues (I335-A345) within β-strand A in EC4, which we denote as “δ-insertion”, is also unique and not seen in other clustered protocadherin structures (Fig. 3e and Supplementary Fig. 5). Similarly, a six-residue insertion in the DE loop of EC4 (G392-K397) is found only in PCDH1 sequences (Supplementary Fig. 5 and Supplementary Table 3). These unique features might be relevant for function, as further discussed below.

There are three cysteine residues found in the EC1-4 sequence, two of which are involved in an intra-molecular disulfide bond (C66-C75) as discussed above. The third one, C375, is partially buried in both structures (from bacterial and mammalian expressed proteins). This suggests that potential disulfide-mediated dimers observed in refolded PCDH1 EC1-4 might be rare or artefactual.

The PCDH1 EC1-4mc structure had regions in which electron density could only be explained by the presence of glycosylation. Residues N89, N248 and N346 are predicted to be N-linked glycosylation sites and sugars can be observed at N248 in the PCDH1 EC1-4mc structure. C-linked glycosylation has been predicted for W186 but is not observed in our structure, while O-linked glycosylation has not been predicted but it is observed at S430 (Supplementary Fig. 3b). The role of these glycosylation sites in PCDH1 function is unclear.

### Antiparallel interfaces in the crystal structure of PCDH1 EC1-4 suggest adhesive mechanisms

Relevant adhesive interfaces have been observed in crystal structures of classical cadherins, of the protocadherin-15 and cadherin-23 heterophilic complex, and of clustered and non-clustered protocadherins^18,39,40,42,43,46,47^. The crystal structures of PCDH1 EC1-4 obtained from both bacterial and mammalian cells reveal the same two plausible interfaces of homophilic interaction (Fig. 4). The first homophilic adhesion interface (PCDH1-I1) involves two PCDH1 EC1-4 molecules having complete antiparallel overlap. In this arrangement, EC1 of one molecule interacts with EC4 of the second molecule (EC1:EC4), EC2 with EC3 (EC2:EC3), EC3 with EC2 (EC3:EC2) and EC4 with EC1 (EC4:EC1) (Fig. 4a-b). The second possible interface of adhesion (PCDH1-I2) involved the opposite side of PCDH1 EC1-4, having antiparallel overlap of EC2 of one molecule with EC4 of the other (EC2:EC4), EC3 with EC3 (EC3:EC3), EC4 with EC2 (EC4:EC2) and potential EC1:EC5 contacts (with EC5 not seen in the structure; Fig. 4c-d). These two possible interfaces were analyzed and tested to determine which one mediates adhesion *in vitro* and potentially *in vivo*.

**Figure 4.**
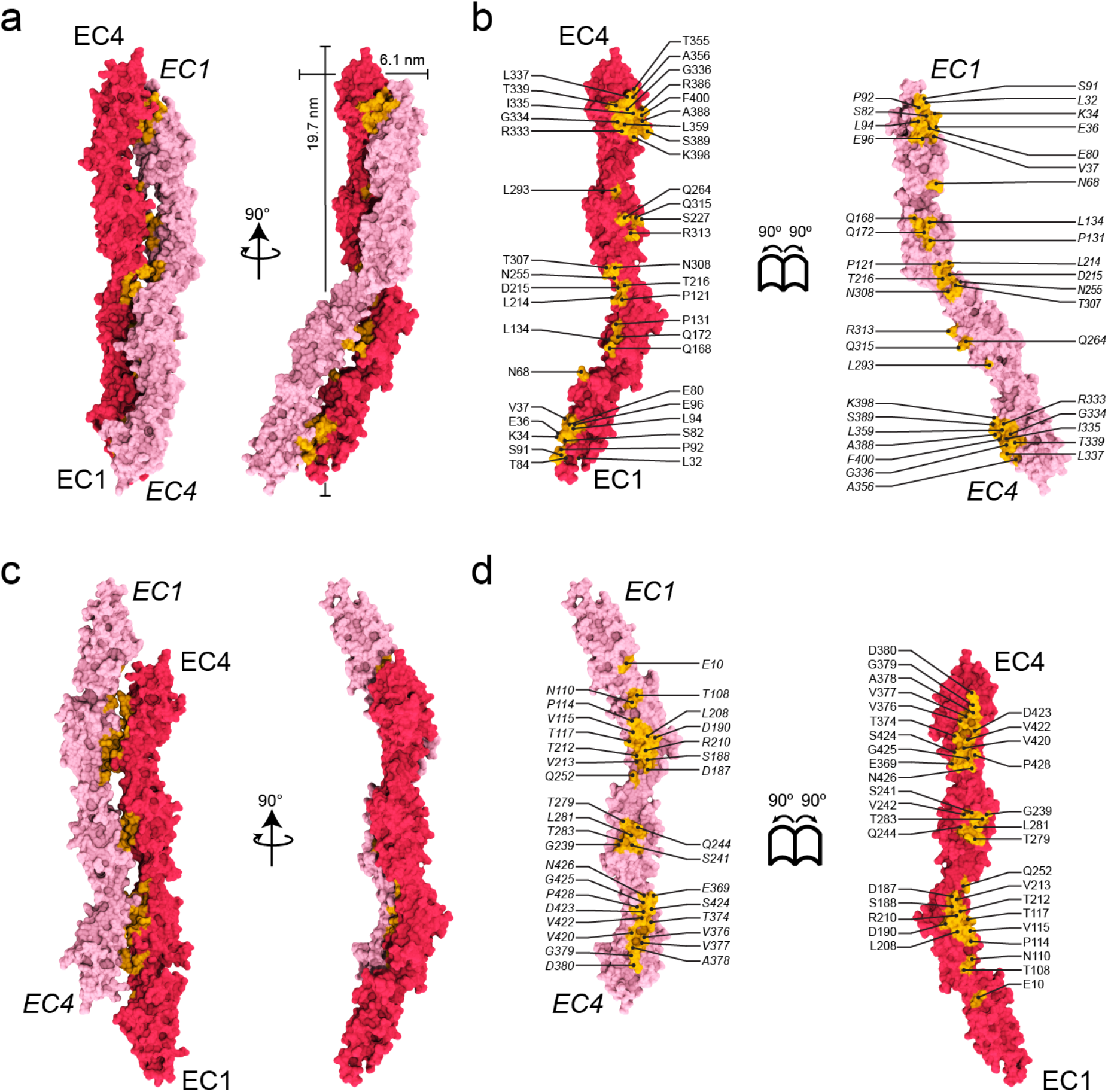
Crystallographic antiparallel interfaces of PCDH1. **(a)** Molecular surface representation of two PCDH1 EC1-4 molecules in the crystal with the interaction interface formed by fully overlapped and antiparallel EC1-4 repeats (PCDH1-I1). Two perpendicular views are shown. **(b)** Interaction surface exposed with interfacing residues listed and shown in yellow. **(c)** Molecular surface representation of two PCDH1 EC1-4 molecules in an alternate crystallographic arrangement in which the interaction interface would be formed by fully overlapped and antiparallel EC1-5 repeats (PCDH1-I2). Two perpendicular views are shown. **(d)** Interaction surface exposed with interfacing residues listed and shown in yellow.

Analyses of the interaction interfaces with the Protein, Interfaces, Surfaces and Assemblies (PISA)^48^ server revealed large interface areas of 1,539 Å^2^ for PCDH1-I1 (385 Å^2^ per repeat; 486 Å^2^ for EC1:EC4 and EC4:EC1, 231 Å^2^ for EC2:EC3 and EC3:EC2; Table 2) and 1,270 Å^2^ for PCDH1-I2 (423 Å^2^ per repeat; 461 Å^2^ for EC2:EC4 and EC4:EC2, 190 Å^2^ for EC3:EC3) respectively. Both interface areas are greater than 856 Å^2^, an empirical cut-off to distinguish biological interfaces from crystal contacts. Therefore both are unlikely to be crystal packing artifacts^49^. Interestingly, the interface area per EC repeat is larger for PCDH1-I2.

**Table 2.**
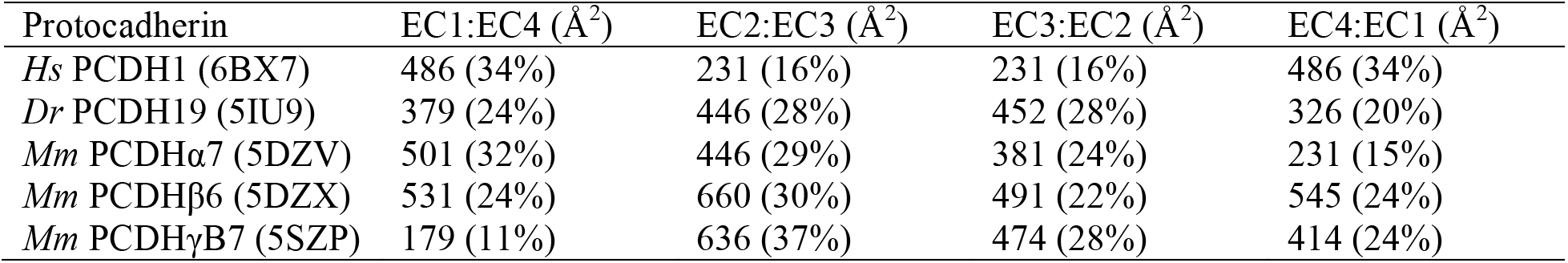
Interface areas per EC repeat for PCDH1 (δ1 protocadherin), PCDH19 (δ2 protocadherin), PCDHα7, PCDHβ6, and PCDHγB7 (clustered protocadherins). The percentage of total interface areas that they cover are in brackets.

| Protocadherin | EC1:EC4 ($\text{\AA}^2$) | EC2:EC3 ($\text{\AA}^2$) | EC3:EC2 ($\text{\AA}^2$) | EC4:EC1 ($\text{\AA}^2$) |
| --- | --- | --- | --- | --- |
| *Hs* PCDH1 (6BX7) | 486 (34%) | 231 (16%) | 231 (16%) | 486 (34%) |
| *Dr* PCDH19 (5IU9) | 379 (24%) | 446 (28%) | 452 (28%) | 326 (20%) |
| *Mm* PCDHα7 (5DZV) | 501 (32%) | 446 (29%) | 381 (24%) | 231 (15%) |
| *Mm* PCDHβ6 (5DZX) | 531 (24%) | 660 (30%) | 491 (22%) | 545 (24%) |
| *Mm* PCDHγB7 (5SZP) | 179 (11%) | 636 (37%) | 474 (28%) | 414 (24%) |

Glycosylation sites, if present at the interface, may interfere with binding and thus reveal non-physiological interfaces as in the case of VE-cadherin^50^. Analyses of the N- and C-linked predicted glycosylation sites showed that residues N89 and N346 are not present in either of the interfaces. In contrast, residues N248 and W186 are near the PCDH1-I2 interface. The observed O-linked glycosylated residue S430 is also near the PCDH1-I2 interface. However, sugars observed at N248 and S430 did not interfere with this interface in the crystal.

Similarly, intermolecular disulfide bonds could interfere with binding thus favoring one interface over the other. Cysteine C375 in EC4 is located near the PCDH1-I2 interface (EC4:EC2), suggesting that an intermolecular disulfide bond at this location would block this interface and favor PCDH1-I1. Yet involvement of C375 in disulfide bond formation is not required for adhesion and might be irrelevant *in vivo*.

Next, we designed mutations predicted to break each interface as determined from an analysis of the structures. A residue was considered to be at the interface if its buried surface area was at least 20% of their accessible surface area according to PISA. Residues K398 in the PCDH1-I1 interface and V115 in the PCDH1-I2 interface met this criterion and were selected for mutagenesis. For the PCDH1-I1 interface, the K398 residue was found to form a salt bridge with E80 of the adjacent molecule. Therefore, we introduced the K398E mutation to prevent the interaction and the formation of the PCDH1-I1 interface. For the PCDH1-I2 interface, V115 was found to form hydrophobic interactions with V420, and therefore we introduced the V115R mutation to disrupt this interface (Fig. 5a,d).

**Figure 5.**
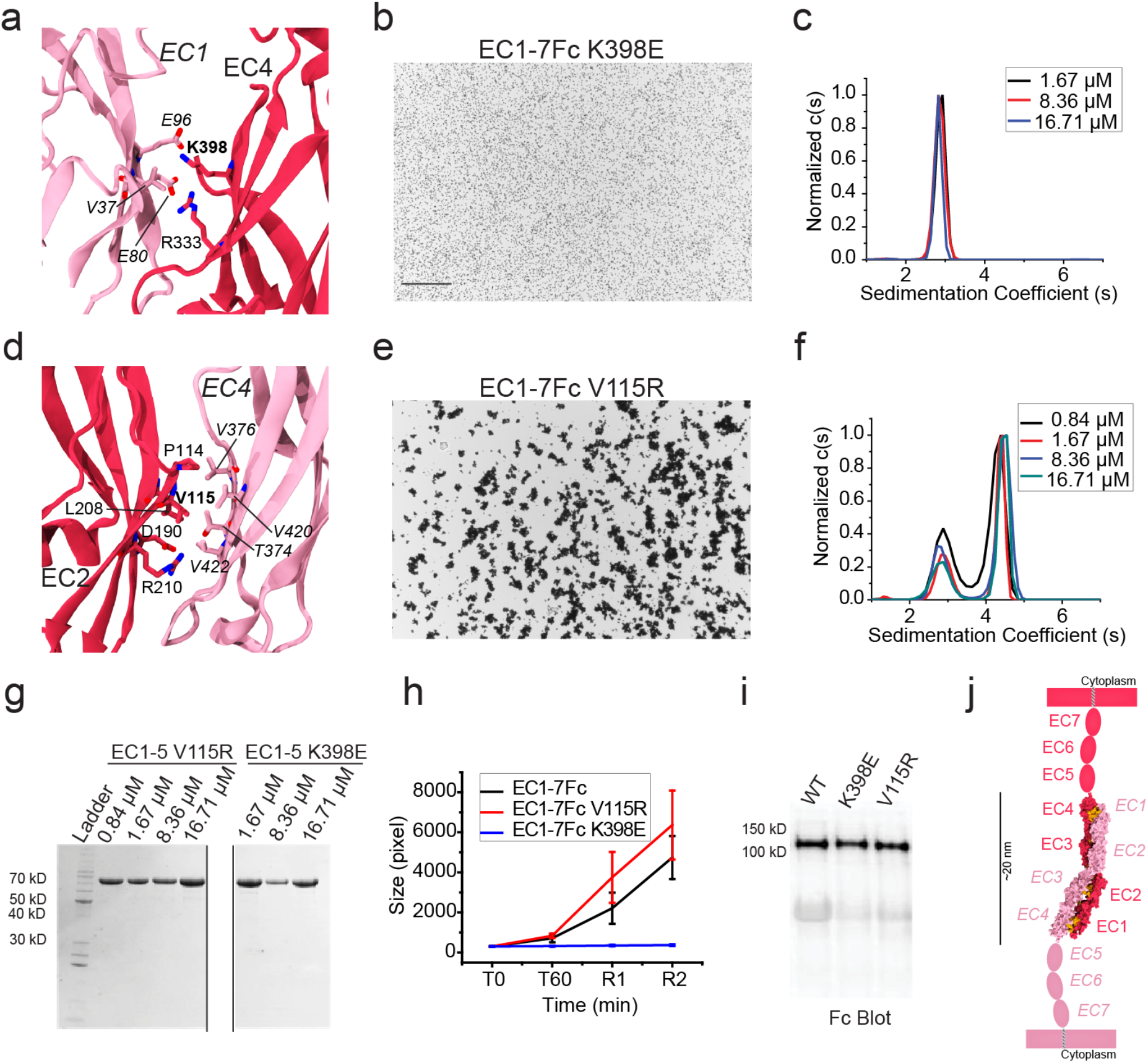
Testing of crystallographic PCDH1 antiparallel interfaces. **(a)** Detail of the PCDH1-I1 interface (EC1:EC4). Residue K398E interacting with E80 is highlighted. **(b)** Protein G beads coated with PCDH1 EC1-7Fc K398E mutant protein and imaged after incubation for 1 hr followed by rocking for 2 min in the presence of calcium. Bar – 500 μm. No aggregation is observed. **(c)** AUC of bacterially produced PCDH1 EC1-5 C375S C548S K398E shows only a monomeric peak for a range of concentrations between 1.67 μM and 16.71 μM. **(d)** Detail of the PCDH1-I2 interface (EC1:EC5). Residue V115 interacting with V420 is highlighted. **(e)** Protein G beads coated with PCDH1 EC1-7Fc V115R mutant protein and imaged after incubation for 1 hr followed by rocking for 2 min in the presence of calcium. Bar – 500 μm. Aggregation is observed. **(f)** AUC of bacterially produced PCDH1 EC1-5 C375S C548S V115R shows both dimeric and monomeric peaks for a range of concentrations between 0.84 μM and 16.71 μM. **(g)** SDS-PAGE shows samples for PCDH1 EC1-5 C375S C548S V115R and PCDH1 EC1-5 C375S C548S K398E after AUC. Different amounts of samples were loaded while running the gel. **(h)** Mean aggregate size for wild type PCDH1 EC1-7Fc and mutants PCDH1 EC1-7Fc K398E, PCDH1 EC1-7Fc V115R at T0 (*t* = 0 min), after 1 hr of incubation, T60 (*t* = 60 min) followed by rocking for 1 min (R1) and 2 min (R2). Error bars are standard error of the mean (*n* = 4 for mutants, *n* = 3 for wild type). **(i)** Western blot shows efficient expression of wild type PCDH1 EC1-7Fc and mutants PCDH1 EC1-7Fc K398E and PCDH1 EC1-7Fc V115R. **(j)** Schematic of *trans* antiparallel binding of PCDH1 involving overlap of repeats EC1 to EC4.

Bead aggregation assays of PCDH1 EC1-7Fc K398E revealed no aggregation of beads in the presence of calcium (Fig. 5b,h,i). Consistent with these results, sedimentation velocity AUC experiments with protein fragments produced in bacteria and with mutated cysteine residues to avoid confounding effects showed monomeric states in solution for the K398E mutant (PCDH1 EC1-4 C375S K398E and PCDH1 EC1-5 C375S C548S K398E; Fig. 5c,g and Supplementary Fig. 6a-b). In contrast, binding assays of PCDH1 EC1-7Fc V115R revealed aggregation of beads in the presence of calcium and AUC experiments showed both monomeric and dimeric states in solution for this mutant (PCDH1 EC1-5 C375S C548S V115R) (Fig. 5e-i). Melting experiments of the cysteine mutated PCDH1 EC1-5 C375S C548S, EC1-5 C375S C548S K398E and EC1-5 C375S C548S V115R showed that their stability is comparable (Supplementary Fig. 7). These results indicate that the PCDH1-I1 interface mediates homodimerization in solution and is responsible for bead aggregation *in vitro*.

### Model for homophilic interaction of PCDH1 and other δ1 protocadherins

Mutagenesis coupled with binding assays and AUC experiments of the wild type and mutant protein fragments strongly suggest that PCDH1-I1 is the interface that mediates homophilic interaction. A schematic of this mode of interaction is shown in Fig. 5j. This *trans* antiparallel dimer interface suggests a mechanism of adhesion involving EC repeats 1 to 4. This mechanism of interaction is different from the tryptophan exchange mode between EC1 repeats of classical cadherins^36^ and the “handshake” mechanism of interaction between protocadherin-15 and cadherin-23 involving EC1 and EC2^39^. It is similar to the “forearm handshake” mechanism of interaction of the δ2 protocadherins and of the clustered protocadherins involving EC1 to EC4^42,43,45^. Although the overall mode of homophilic adhesion might be the same for δ1, δ2, and clustered protocadherins, there are important differences. The EC1:EC4 contacts are greater in the δ1-protocadherin PCDH1, covering almost 64% of the total interface area (486 Å^2^ for EC1:EC4 and 231 Å^2^ for EC2:EC3). In contrast, both PCDH19 (δ2 protocadherin) and clustered protocadherins EC2:EC3 interface areas are much larger than their EC1:EC4 contacts^42,43^ (Table 2).

An analysis of sequence conservation across species of interfacial residues revealed that residues at EC1:EC4 contacts are more conserved than the ones in EC2:EC3 sites (Fig. 6a-b and Supplementary Figs 8 and 9). About 43% of the interfacial area involves hydrophobic residues, ~23% are charged, and ~34% are hydrophilic residues (Supplementary Fig. 10). The EC2:EC3 contacts are more amphiphilic with ~42% hydrophobic, ~17% charged, and ~42% hydrophilic contacts as compared to EC1:EC4 contacts which are ~43% hydrophobic, ~26% charged and ~30% hydrophilic. All salt-bridge interactions are present in the EC1:EC4 interface. Thus, the size and charged nature of the PCDH1 EC1:EC4 interface suggest that it is important for both strength and specificity of PCDH1 homophilic adhesion.

**Figure 6.**
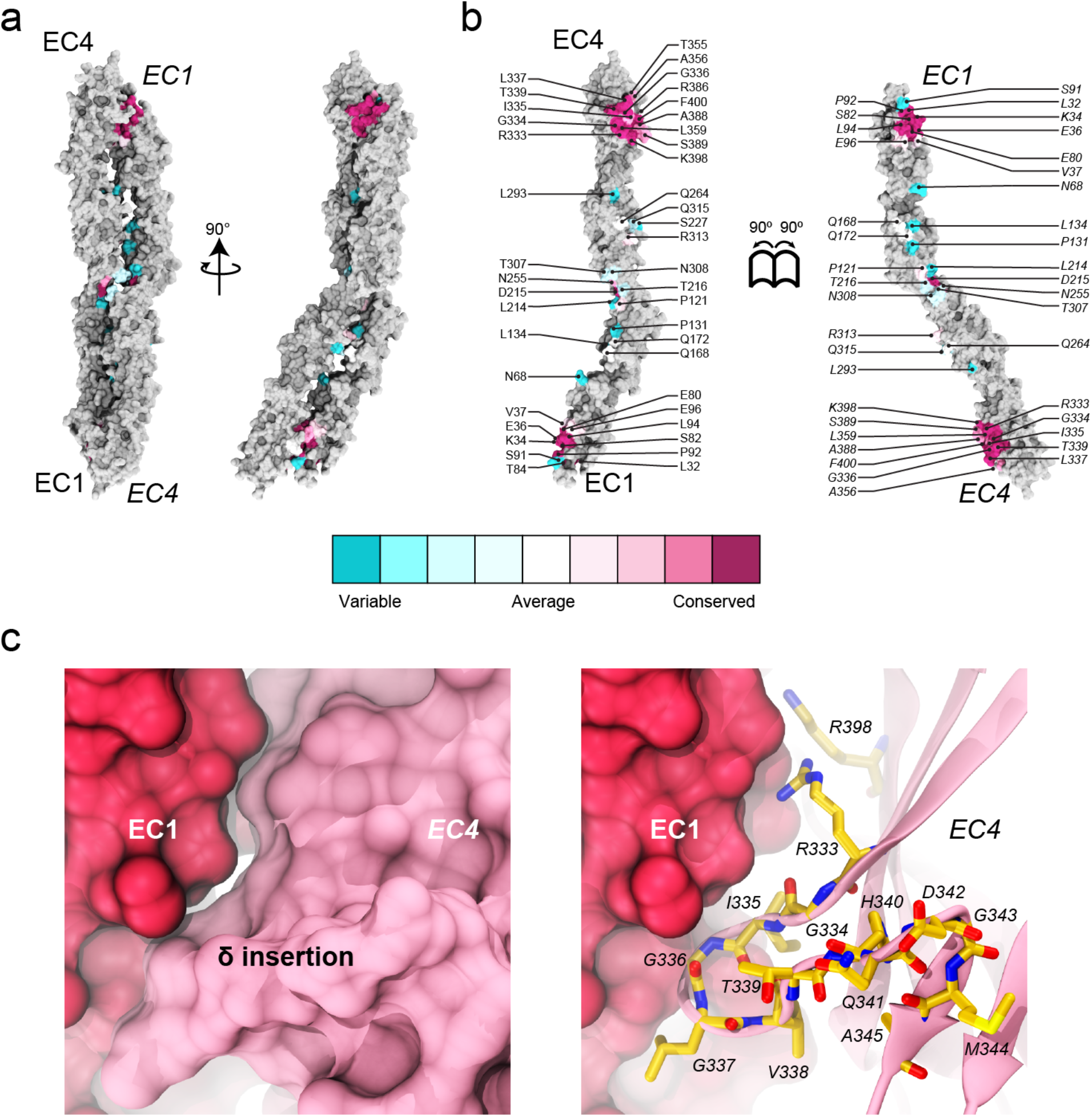
Conservation of residues at the crystallographic PCDH1-I1 interface. **(a)** Molecular surface representation of PCDH1-I1 antiparallel dimer. Two perpendicular views are shown. **(b)** Interaction surface exposed with interfacing residues listed and colored according to sequence conservation among 87 species (right). The accession numbers for the sequences of 87 species are given in Supplementary Table 1. **(c)** Detail of the δ insertion in EC4 shown in molecular surface representation (left) and in ribbon with relevant residues as sticks (right).

An interesting structural aspect of the PCDH1 EC1:EC4 interface is the involvement of the δ insertion within β-strand A of EC4. This eleven-residue insertion contributes four residues that directly contact EC1 of the opposite protomer (Fig. 6c). Similar insertions are found in other mouse and human δ1 protocadherins and the δ2 protocadherin-8, but not in clustered protocadherins. The δ insertion may modulate the affinity of the interaction by providing additional contacts specific to each type of δ protocadherin.

Given the sequence similarity among δ1 protocadherins (~50%) (Supplementary Figs 11-14), the mode of interaction is likely to be the same for all other δ1 family members – PCDH7, PCDH9, and PCDH11X/Y. In support of this model, a very recent study showed that PCDH9 also uses its EC1-4 repeats to mediate homophilic adhesion^51^. A general mode of interaction in which all δ1 protocadherins, with 7 EC repeats, use the same EC1-4 interface used by the δ2 and clustered protocadherins (6 EC repeats) is appealing, but rises the question of how the longer δ1 protocadherins span the intercellular space when compared to the shorter δ2 and clustered protocadherins complexes. Interestingly, PCDH1 EC1-4 is more curved than the clustered protocadherins and PCDH19 EC1-4 fragments (Supplementary Fig. 15 and Videos 1 and 2). Comparison of the PCDH1-I1 complex with those formed by δ2 and clustered protocadherins also shows that the PCDH1 arrangement of protomers is more twisted, yet significant tilting would still be required to span the same intercellular space that the other shorter members of the family can cover using the same EC1-4 binding mechanism.

## DISCUSSION AND CONCLUSIONS

Non-classical cadherins are part of multiple and diverse structures in multicellular organisms, including adhesive bonds important for morphogenesis, sensory perception, and neuronal connectivity^11–13,36^. Structural features of these cadherins are key determinants of their function. For instance, long non-classical cadherins such as protocadherin-24 (9 ECs), protocadherin-15 (11 ECs), and cadherin-23 (27 ECs), are part of long protein filaments essential for enterocyte brush border morphogenesis and inner-ear mechanotransduction^11,52^. In contrast, the shorter clustered protocadherins, with 6 EC repeats, have been implicated in neuronal self-recognition and avoidance, and somewhat paradoxically have been shown to form weak and strictly homophilic *trans* adhesive bonds^13^.

In the context of brain wiring, δ protocadherins are unique. First, unlike the clustered protocadherins, the cytoplasmic domains of δ1 and δ2 protocadherins have distinct sequence motifs that suggest a more diverse and versatile role in cellular signaling^19^. Second, a unique δ insertion in EC4 hints at a specialized role for this repeat in δ protocadherins. Last, the length of δ1 protocadherins’ extracellular domains, with 7 EC repeats, suggest a distinct set of functional roles yet to be determined. Given these differences and their differential expression in the brain^20,26,27,53,54^, an important question about δ protocadherins’ function pertains to whether they are involved in neuronal adhesion and circuitry specificity or rather mediate neuronal self-recognition and avoidance as proposed for the clustered protocadherins.

Functional studies of PCDH19 indicate that this δ2 protocadherin is required for the formation of neuronal columns in the fish optic tectum, thus implying an adhesive role^55^. Structural studies, validated with analyses of epilepsy-causing mutations, indicate that PCDH19 uses an extensive but weak antiparallel EC1-4 adhesive interface ideally suited to encode specificity^42^. Clustered protocadherins use the same antiparallel EC1-4 interface^40,43–45^, and sequence and evolutionary analyses suggest that at least some subgroups of clustered protocadherins use their EC1:EC4 contacts to modulate the bond affinity and their EC2:EC3 interactions to determine specificity^45^.

Interestingly, our structures of PCDH1 and associated binding assays indicate that this δ1 protocadherin uses an antiparallel EC1-4 interface that is similar to that used by clustered and δ2 protocadherins, yet interactions at the EC2:EC3 contacts are minimal, and salt-bridges possibly encoding for specificity are at the much larger EC1:EC4 interface. Binding assays of PCDH9, another δ1 protocadherin, suggest that this protein also uses an antiparallel EC1-4 interface, with the EC1:EC4 contact playing a more important role than EC2:EC3 interactions in bead aggregation^51^. Taken together, these results indicate that δ1 protocadherins use a unique and enlarged EC1:EC4 contact both to strengthen the adhesive bond and to determine its specificity. Whether δ1 protocadherins use this bond to trigger a repulsive signal mediating neuronal self-recognition, to form true adhesive and specific intercellular contacts, or both depending on molecular and cellular context, is still unclear.

The versatility of PCDH1 in mediating adhesion and signaling might be enhanced by the existence of various isoforms that modify its extracellular and cytoplasmic domains^56^. The possible existence of a PCDH1 isoform with a shorter extracellular domain starting at EC4 (NM_001278615.1 / NP_001265544.1) indicates that a pseudo-heterophilic bond could be formed and be mediated by the EC1:EC4 interface alone. Surface expression levels of this isoform could modulate the strength of the adhesion between two cells expressing PCDH1.

The discovery of an antiparallel EC1-4 interface for PCDH1 and δ1 protocadherins with 7 ECs poses an interesting set of questions regarding the arrangement of this adhesive complex. The intercellular space occupied by classical cadherins (5 ECs interacting through EC1s in a curved and tilted arrangement, ~25 nm)^47^ or by clustered and δ2 protocadherins (6 ECs with an EC1-4 overlap, <40 nm)^42,44,45^ might be too short to accommodate a straight δ1 protocadherin complex that uses the EC1-4 antiparallel interface and that has 7 ECs per protomer (<50 nm). While PCDH1 is curved, and the complex is twisted, this might shorten the end-to-end distance of the bond only by a few nanometers, not enough to accommodate two additional ECs (~10 nm) at an intercellular space that could be dominated by shorter clustered and δ2 protocadherins. Tilting of the complex might solve this conundrum. It is also possible that the longer δ1 protocadherins are used to explore and form initial transient cell-cell contacts while other shorter cadherins and protocadherins are recruited to enable more permanent adhesion. On the contrary, a longer and straight δ1 complex might prevent adhesion mediated by shorter and tilted classical complexes. Determining how protocadherins orient with respect to the membrane plane, as recently explored for protocadherin-15^57^, will be critical to elucidate their function.

The similarities in the overall mode of adhesion for δ1, δ2 and clustered protocadherins suggest that they may have a common ancestry^58^. It is thus tempting to speculate that δ protocadherins may engage in *cis* interactions as the clustered protocadherins do using their EC5 and EC6 repeats^41,59^. Whether EC7 in δ1 protocadherins is also involved in this type of contacts is unknown. Interestingly, PCDH19, a δ2 protocadherin, interacts with the classical N-cadherin^42,60,61^. Perhaps PCDH1 interacts with E-cadherin in a similar fashion, as they co-localize in airway epithelial cells. Since the protocadherins are known to mediate weak interactions, *cis* dimerization with classical members of the family may help in forming more robust adhesion complexes.

PCDH1 is also expressed in several non-neuronal tissues, including the cochlea, skin keratinocytes, lungs, and airway epithelia, all of which are subjected to mechanical stimuli^25,28,31,62–64^. The twisted adhesive bond of PCDH1 may provide some elasticity to these tissues, and its length might allow for additional stretching of cell-cell contacts without complete rupture of the adhesive contact. In addition, PCDH1 is known to help in forming the airway epithelial barrier by mediating adhesion between epithelial cells, the dysfunction of which leads to pathogenesis^31^. PCDH1 expression increases during differentiation of airway and bronchial epithelial cells, and it has been localized to cell-cell contact sites along with E-cadherin^31^. Our structural model of PCDH1 homophilic adhesion partially reveals how the airway epithelial barrier is formed, and suggests mutation sites that can be used to disrupt PCDH1 adhesion, which should help to elucidate the roles played by PCDH1’s extracellular and cytoplasmic domains *in vivo*.

The structure of PCDH1 EC1-4 and the homophilic binding mechanism presented here also have important implications in relation to New World hantaviruses infectivity. PCDH1 EC1 has been recently shown to be critical for cellular attachment and entry of these viruses^32^, yet molecular details are still unclear. It is possible that the hantavirus glycoproteins interact with PCDH1 EC1 and disrupt homophilic binding by preventing contacts with the partnering PCDH1 EC4. In this scenario, PCDH1 EC4 could be used and engineered to block interactions with the virus. On the other hand, native PCDH1 EC1 interactions with hantavirus glycoproteins may involve residues not engaged in homophilic binding, in which case PCDH1 EC1 itself could be used to develop hantavirus blockers. In either case, the molecular details provided by our PCDH1 EC1-4 structure may guide drug discovery strategies aimed at reducing or blocking hantavirus infections.

## METHODS

### Cloning and Mutagenesis

Sequences enconding for human PCDH1 repeats EC1-4 and EC1-5 were subcloned using the NdeI and XhoI sites of the pET21a vector for bacterial expression (without signal peptide), and using the XhoI and BamHI sites of the CMV:N1-Fc and CMV:N1-His vector^42^ for mammalian expression. Constructs encoding for truncated versions of PCDH1 (PCDH1EC1-7, PCDH1EC1-6, PCDH1EC1-5, PCDH1EC1-4, PCDH1EC1-3, PCDH1EC1-2, PCDH1EC2-7) were created by PCR subcloning of the native signal peptide and appropriate EC domains into CMV:N1-Fc. Truncation boundaries for EC repeats are as in Supplementary Fig. 4. Mutations were created in both the bacterial and the mammalian expression constructs by site-directed mutagenesis using the Quikchange Lightning Site-directed Mutagenesis Kit (Agilent). All constructs were sequence verified.

### Bacterial Expression, Purification, and Refolding

PCDH1 fragments were expressed in BL21 CodonPlus (DE3)-RIPL cells (Stratagene) that were cultured in LB (EC3-4, EC1-4 and EC1-5 WT and mutants), induced at OD_600_ ~ 0.6 with 200 μM IPTG, and grown at 30 °C for ~18 hr. Cells were lysed by sonication in denaturing buffer (20 mM TrisHCl [pH 7.5], 6 M guanidine hydrochloride, 10 mM CaCl_2_ and 20 mM imidazole). The cleared lysates were loaded onto Ni-Sepharose (GE Healthcare) and eluted with denaturing buffer supplemented with 500 mM imidazole. PCDH1 fragments for crystallography were refolded by overnight dialysis against 20 mM TrisHCl [pH 8], 150 mM KCl, 50 mM NaCl, 400 mM arginine, 2 mM CaCl_2_, and 2 mM DTT using MWCO 2000 membranes at 4 °C. Refolded proteins were further purified by size exclusion chromatography on a Superdex200 column (GE Healthcare) in 20 mM TrisHCl [pH 8.0], 150 mM KCl, 50 mM NaCl, 2 mM CaCl_2_ and 2 mM DTT. PCDH1 fragments used for analytical ultracentrifugation experiments were refolded and purified using the same buffers without DTT.

### Mammalian Expression and Purification

The PCDH1 EC1-4His fragment was expressed in Expi293^TM^ cells (ThermoFischer Scientific). Transfection was done with 30 μg of plasmid and ExpiFectamine^TM^ 293. Media was collected on the fourth day after transfection by pelleting cells and collecting the supernatant to which 10 mM CaCl_2_ was added. The mix was dialysed overnight against 20 mM TrisHCl [pH 8.0], 150 mM KCl, 50 mM NaCl, and 10 mM CaCl_2_ using MWCO 2000 membranes. The PCDH1 EC1-4His fragment was then purified by nickel affinity chromatography under native conditions with buffer containing 20 mM TrisHCl [pH 8.0], 300 mM NaCl, 10 mM CaCl_2_ and 20 mM imidazole. The protein was eluted with the same buffer containing 500 mM imidazole. It was further purified by size exclusion chromatography on a Superdex200 column (GE Healthcare) in 20 mM TrisHCl [pH 8.0], 150 mM KCl, 50 mM NaCl, and 2 mM CaCl_2_.

### Analytical Ultracentrifugation

Sedimentation velocity experiments were performed in a ProteomeLab™XL-I analytical ultracentrifuge (Beckman Coulter). Samples were prepared by concentrating purified proteins from size exclusion chromatography using Sartorious Vivaspin 20 concentrators (MWCO 10 kD). Protein concentrations were measured using a Nanodrop 2000c spectrophotometer (Thermo Scientific). Buffer containing 20 mM TrisHCl [pH 8.0], 150 mM KCl, 50 mM NaCl and 2 mM CaCl_2_ was used as blank. All samples were run at 50,000 rpm at 25 °C and detection was by UV at 230 nm for samples with concentration < 8 μM and at 280 nm for others. Solvent density was assumed to be 1 g/cc and the partial specific volume of solute 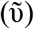 was set to 0.73 cc/g. Data was analyzed using SEDFIT^65,66^.

### Crystallization, Data Collection, and Structure Determination

Crystals of PCDH1 EC1-4 fragments from bacterial and mammalian expression systems were grown by vapor diffusion at 4 °C by mixing equal volumes of protein (~ 6 mg/ml) and reservoir solution (0.1 M NaHEPES [pH 7.5], 0.2 M CaCl_2_, 15% PEG 400, 15% Glycerol for PCDH1 EC1-4bc; 0.05 M TrisHCl [pH 8.5], 0.1 M MgCl_2_, 30% PEG550 MME for PCDH1 EC1-4mc). Crystals of PCDH1 EC3-4 fragments from bacterial expression systems were also grown by vapor diffusion at 4 °C by mixing equal volumes of protein (~ 8.5 mg/ml) and reservoir solution of 0.1 M MES [pH 6.5], 20% PEG 4000, and 0.6 M NaCl (cryo-protected with 25% glycerol). Crystals were cryo-cooled in liquid N_2_. X-ray diffraction data was collected as indicated in Table 1 and processed with HKL2000^67^. The PCDH1 EC1-4bc structure was determined by molecular replacement using an unfinished PCDH1 EC3-4 structure as an initial search model in PHASER^68^. Model building was done with COOT^69^ and restrained TLS refinement was performed with REFMAC5^70^. Likewise, the PCDH1 EC1-4mc structure was determined through molecular replacement using the PCDH1 EC1-4bc structure as the initial search model in PHASER. Data collection and refinement statistics are provided in Table 1. Inspection of 2Fo-Fc electron density maps during initial stages of refinement of the PCDH1 EC1-4bc and PCDH1 EC1-4mc model revealed unassigned densities at the linker regions. Ca^2+^ was present at concentrations of at least 2 mM in the protein solution buffers, so we assigned Ca^2+^ ions in the PCDH1 EC1-4bc model. However, the crystallization buffer for PCDH1 EC1-4mc had 0.1 M MgCl_2_. We tried to assign both Ca^2+^ as well as Mg^2+^ in the linker regions of the PCDH1 EC1-4mc model, but only the assignment of Ca^2+^ ions was compatible with the final 2Fo-Fc map and was supported by an analysis of distances to coordinating atoms and B factor values of the ion and surrounding residue atoms. All molecular images were generated with VMD^71^.

### Bead Aggregation Assays

Bead aggregation assays were modified from those described previously^42,61,72,73^. The PCDH1Fc fusion constructs were transfected into HEK293T cells using calcium-phosphate transfection^74–76^. Briefly, solution A (10 μg of plasmid DNA and 250 mM CaCl_2_) was added drop-wise to solution B (2X HBS) while mildly vortexing, and the final transfection solution was added drop-wise to one 100-mm dish of cultured HEK293T cells (two plates per assay). The next day, cells were rinsed twice with 1X PBS and serum-free media. Cells were allowed to grow in the serum-free media for 2 days before collecting the media containing the secreted Fc fusions. The media was concentrated (Amicon 10 kD concentrators) and incubated with 1.5 μl of protein G Dynabeads (Invitrogen) while rotating at 4 °C for 2 hr. The beads were washed in binding buffer (50 mM TrisHCl [pH 7.5], 100 mM NaCl, 10 mM KCl, and 0.2% BSA) and split into two tubes with either 2 mM EDTA or 2 mM CaCl_2_. Beads were allowed to aggregate in a glass depression slide in a humidified chamber for 60 min without motion, followed by 1 min and then 2 min of rocking (8 oscillations/min). Images were collected upon adding EDTA or CaCl_2_, after 60 min incubation, and after each rocking interval using a microscope (Nikon Eclipse Ti) with a 10x objective. Bead aggregates were quantified using ImageJ software as described previously^61,72^. Briefly, the images were thresholded, the area of the detected aggregate particles was measured in units of pixels, and the average size was calculated. Assays were repeated four times (except for PCDH1 EC1-7Fc and PCDH1 EC1-7Fc C375S C548S with *n* = 3) from separate protein preps and their mean aggregate size (± SEM) at each time point was plotted.

Western blots were performed using part of the media containing the Fc fusion proteins after the bead aggregation assay to confirm expression and secretion of the protein. The media was mixed with sample loading dye, boiled for 5 min and loaded onto SDS-PAGE gels (BioRad) for electrophoresis. Proteins were transferred to PVDF membrane (GE healthcare) and blocked with 5% nonfat milk in TBS with 0.1% Tween-20 before incubating overnight at 4 °C with goat anti-human IgG (1:200 Jackson ImmunoResearch Laboratories). After several washes with 1X TBS, the blot was incubated with mouse anti-goat HRP-conjugated secondary (1:5000, Santa Cruz Biotechnology) for 1 hr at room temperature, washed with 1X TBS and developed with ECL Select Western Blot Detection (GE Healthcare) for chemiluminescent detection (Omega Lum G).

### Differential Scanning Fluorimetry

The bacterially expressed PCDH1 EC1-5 C375S C548S, EC1-5 C375S C548S K398E, and EC1-5 C375S C548S V115R fragments were purified as described above and used for differential scanning fluorimetry (DSF)^77–79^. Experiments were repeated three times using protein at 0.2 mg/ml for all constructs in buffer (20 mM TrisHCl [pH 8.0], 150 mM KCl, 50 mM NaCl, and 2 mM CaCl_2_) mixed with SYPRO Orange dye (Invitrogen). Fluorescent measurements were performed in a CFXConnect RT-PCR instrument (BioRad) using hard-shell 96-well thin wall PCR plates (BioRad) while samples were heated from 10° to 95° in 0.2° steps. Melting temperatures were defined as those recorded when the normalized fluorescence reached 0.5.

### Sequence Analysis

For analysis of PCDH1 residue conservation across species, 87 sequences were obtained from the NCBI protein database and processed manually to include the extracellular domain from EC repeats 1 to 7. The accession numbers for sequences of each of the 87 species is listed in Supplementary Table 1. These PCDH1 sequences were aligned using Clustal X2^80^ and the alignment file was put into ConSurf^81^ to calculate relative conservation of each residue and to categorize the degree of conservation into nine color bins. Sequences for δ1 protocadherins *hs* PCDH1, *mm* PCDH1, *hs* PCDH7, *mm* PCDH7, *hs* PCDH9, *mm* PCDH9, *hs* PCDH11X and *mm* PCDH11X were obtained from the NCBI protein database, processed manually to include only the extracellular domain through the end of EC7, and aligned using the Clustal Omega server^82^. Identity and similarity were calculated using the Sequence Identity And Similarity SIAS server^83^. The accession numbers of the sequences are listed in Supplementary Table 2.

### Prediction of Glycosylation and Glycation Sites

Potential PCDH1 glycosylation sites were predicted using the following servers: NetNGlyc 1.0 (N-glycosylation, GlcNAc-β-Asn), NetOGlyc 4.0 (O-glycosylation, GalNAC-α-Ser/Thr)^84,85^, and NetCGlyc 1.0 (C-glycosylation, Man-α-Trp)^86^.

### Orientation of EC Repeats and Calculation of Azimuthal Angles

Relative orientation of EC repeats was determined using *mm* CDH23 (2WHV) as a reference. Repeat EC1 of *mm* CDH23 was aligned to the *z-*axis in VMD. Then, the N-terminal EC repeat for a pair of ECs of interest was aligned to *mm* CDH23 EC1 using COOT. The principal axis of the C-terminal EC repeat was then calculated using the Orient plugin in VMD. The projection of the principal axis on the *xy*-plane was plotted to identify the azimuthal angle φ. Repeat CDH23 EC2 was used to define φ = 0°.

## ACCESSION NUMBERS

Coordinates for *hs* PCDH1 EC1-4bc and *hs* PCDH1 EC1-4mc have been deposited in the Protein Data Bank with entry codes 6BX7 and 6MGA, respectively.

## Supporting information

## ACKNOWLEDGEMENTS

We thank members of the Jontes and Sotomayor laboratories and Dr. Sharon R. Copper for training, assistance, and discussions. This work was supported in part by the Ohio State University. Use of APS NE-CAT beamlines was supported by NIH (P30 GM124165 & S10 RR029205) and the Department of Energy (DE-AC02-06CH11357) through grants GUP 49774 and 59251. DM is a Pelotonia fellow and MS was an Alfred P. Sloan fellow (FR-2015-65794).

## AUTHOR CONTRIBUTIONS

D.M. did cloning, protein expression and protein purification, and carried out all binding and bead aggregation assays, crystallization trials, differential scanning fluorimetry experiments, and sequence alignments. M.S. trained D.M. and assisted with crystal fishing and cryo-cooling. D.M and M.S. solved crystal structures. M.S. supervised work and assisted with data analysis. D.M. and M.S. prepared figures and wrote the manuscript.

