## Supplementary material for "An Adhesive Interface for the Non-Clustered δ1 Protocadherin-1 Involved in Respiratory Diseases"

December 2018

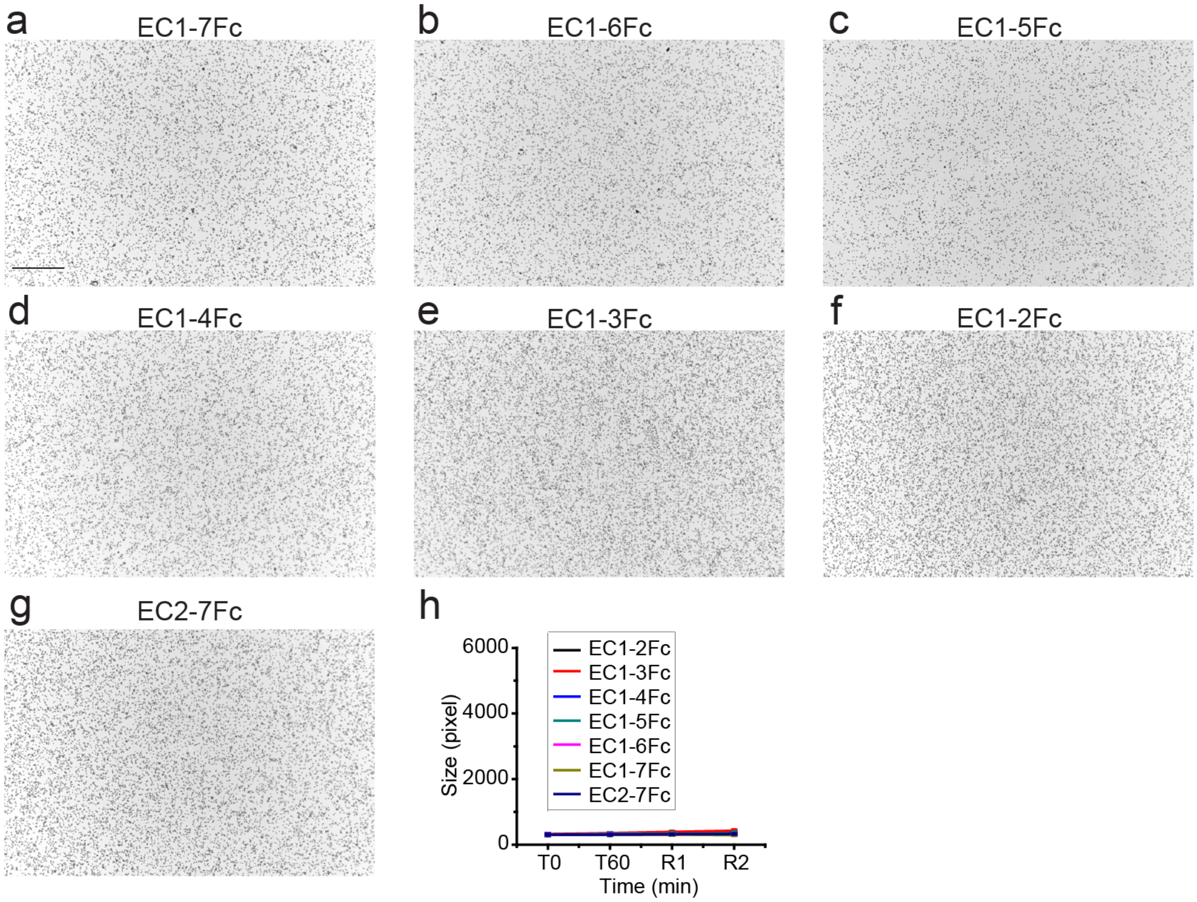


**Supplementary Figure 1. PCDH1 does not mediate adhesion in the absence of calcium (2 mM EDTA). (a-g)** Protein G beads coated with full length **(a)** and truncated versions **(a-g)** of the PCDH1 extracellular domain imaged after incubation for 1 hr followed by rocking for 2 min in the absence of calcium. Bar – 500 µm. **(h)** Mean aggregate size for full length and truncated fragments of PCDH1 at T0 ($t$ = 0 min), after 1 hr of incubation, T60 ($t$ = 60 min) followed by rocking for 1 min (R1) and 2 min (R2). Error bars are standard error of the mean ($n$ = 4 for all constructs except for PCDH1 EC1-7Fc with $n$ = 3).

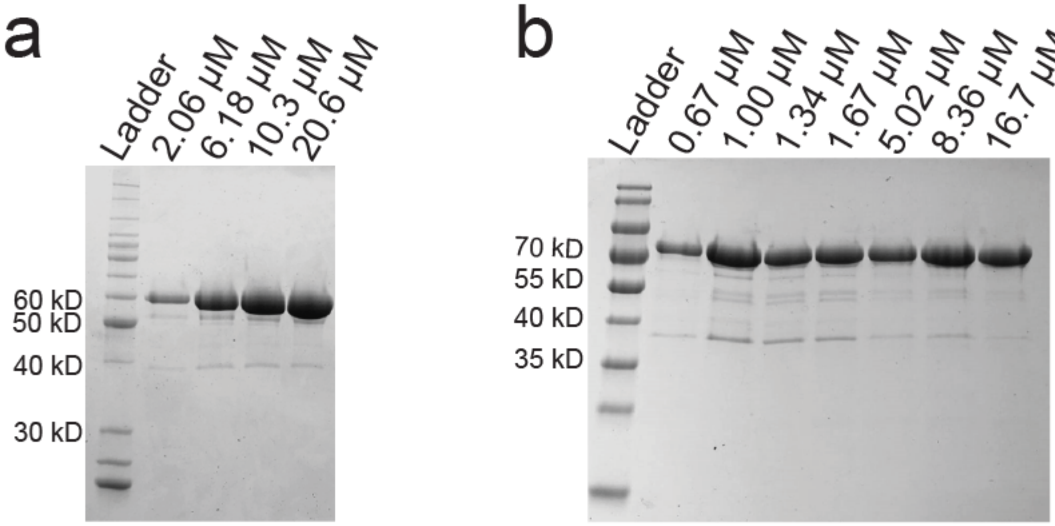


**Supplementary Figure 2. SDS-PAGE analyses of samples after AUC. (a-b)** Gels show that samples for PCDH1 EC1-4 C375S **(a)** and PCDH1 EC1-5 C375S C548S **(b)** were pure and did not degrade after AUC. Samples were not loaded in equal amounts for running the gel.

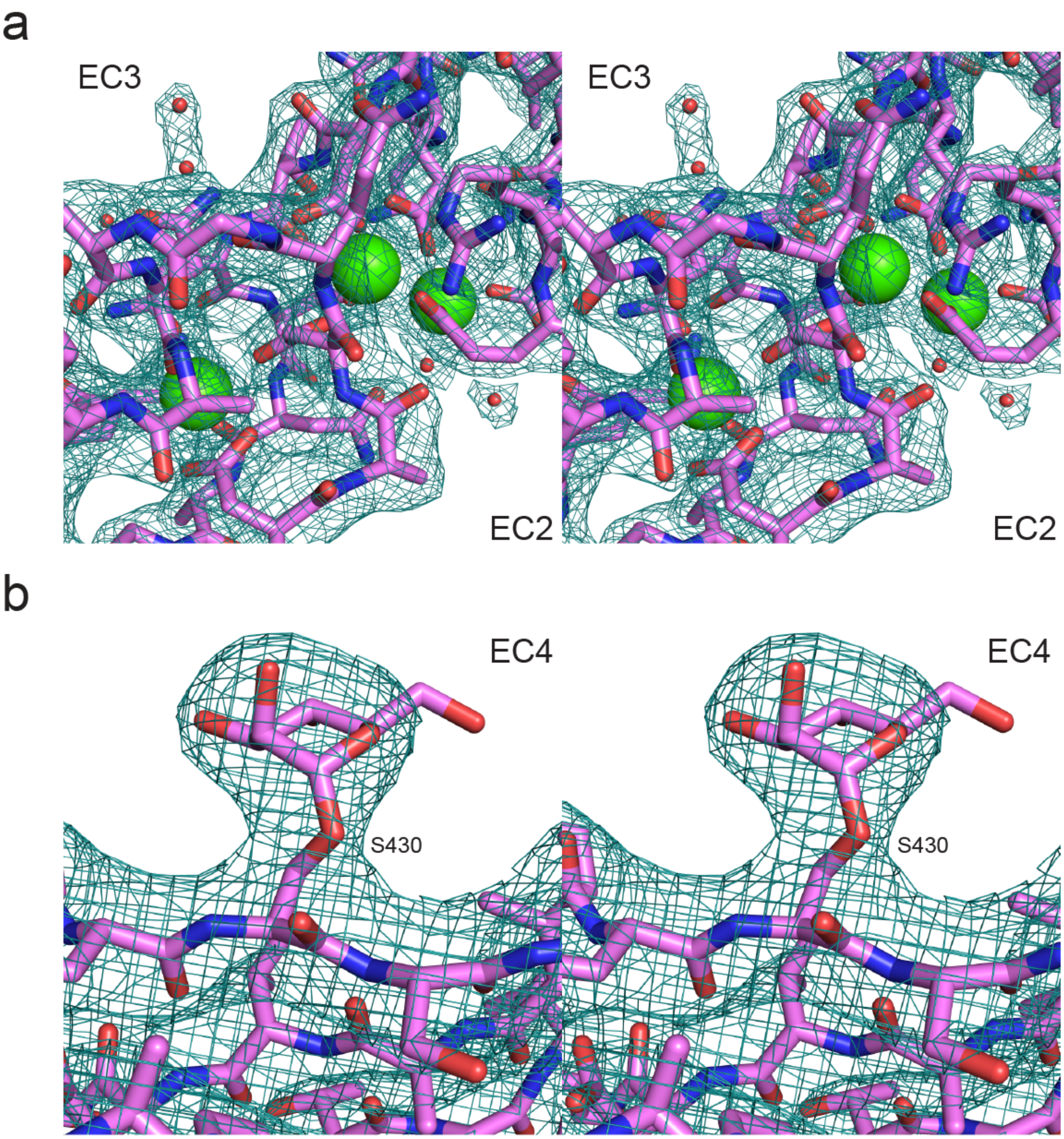


**Supplementary Figure 3. Electron density maps of PCDH1 structures. (a)** Stereo view of the $2F_O$-Fc electron density map (teal mesh) of the EC2-3 linker region in the PCDH1 EC1-4bc structure (2.85 Å; 6BX7) contoured at 2.0 σ. **(b)** Stereo view of the $2F_O$-Fc electron density map (teal mesh) of the mannose sugar molecule in the PCDH1 EC1-4mc structure (3.15 Å; 6MGA) contoured at 1.3 σ.

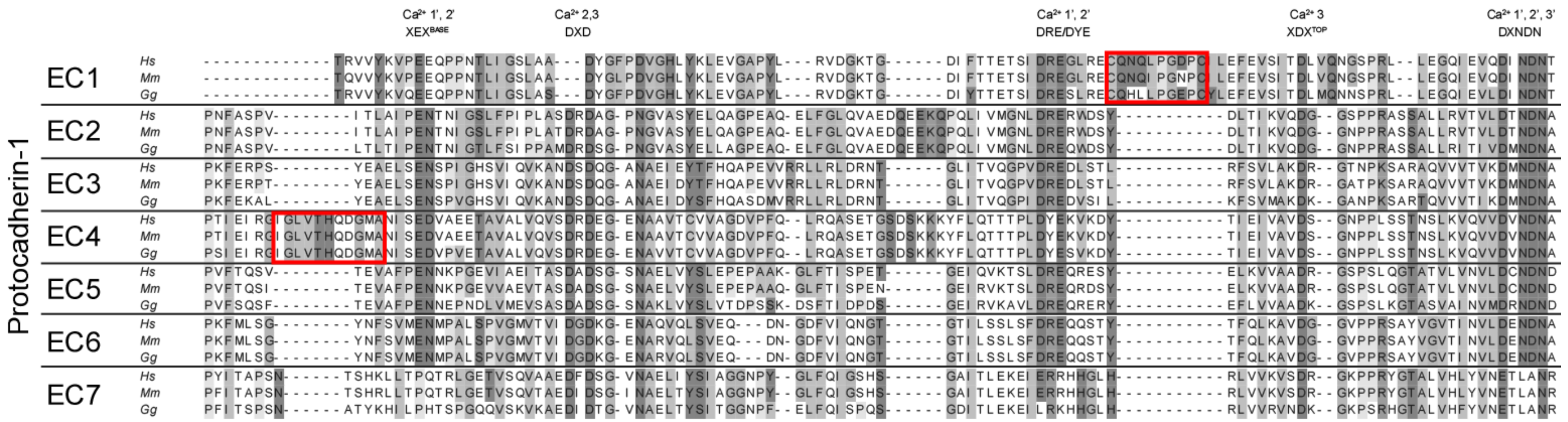


**Supplementary Figure 4. Sequence alignment of human (*hs*), mouse (*mm*), and chicken (*gg*) PCDH1 EC repeats.** All 7 EC repeats for each species are aligned to each other (EC1 to EC7). Conserved calcium-binding motifs are labeled. Red boxes show the cysteine loop in EC1 and the δ insertion in EC4.

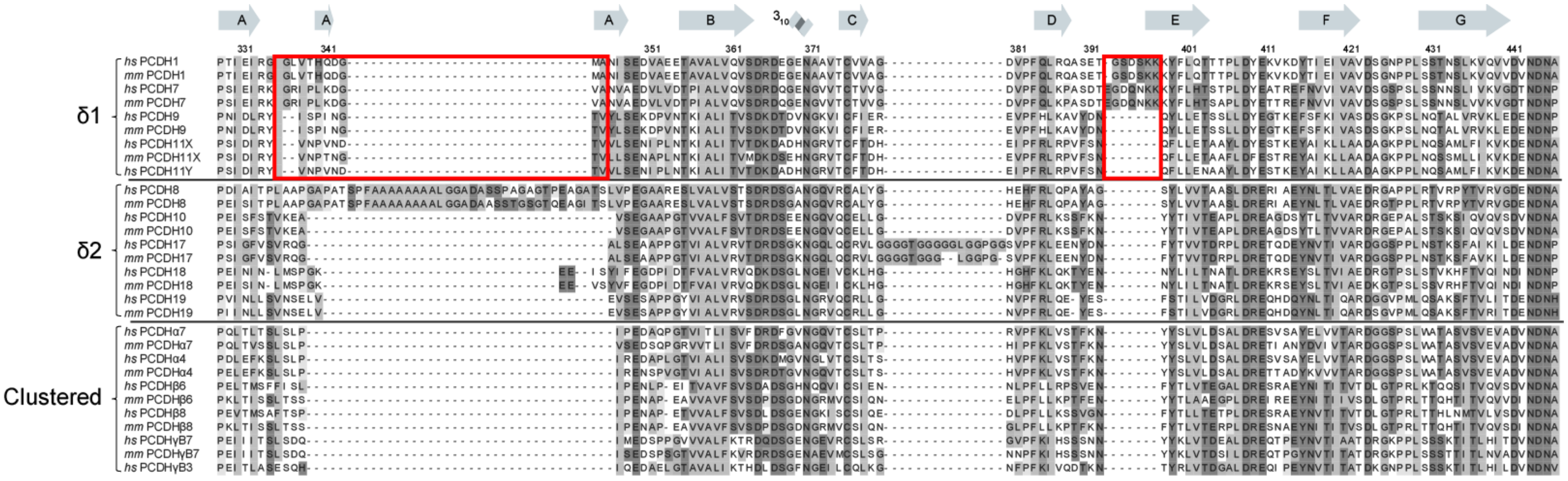


**Supplementary Figure 5. Sequence alignment of repeat EC4 from selected δ and clustered protocadherins.** Secondary structure from *hs* PCDH1 EC4 is indicated on top of the alignment. Red boxes show the δ insertion and a unique DE loop in PCDH1. The accession numbers for the protein sequences used are given in Supplementary Tables 2 and 3.

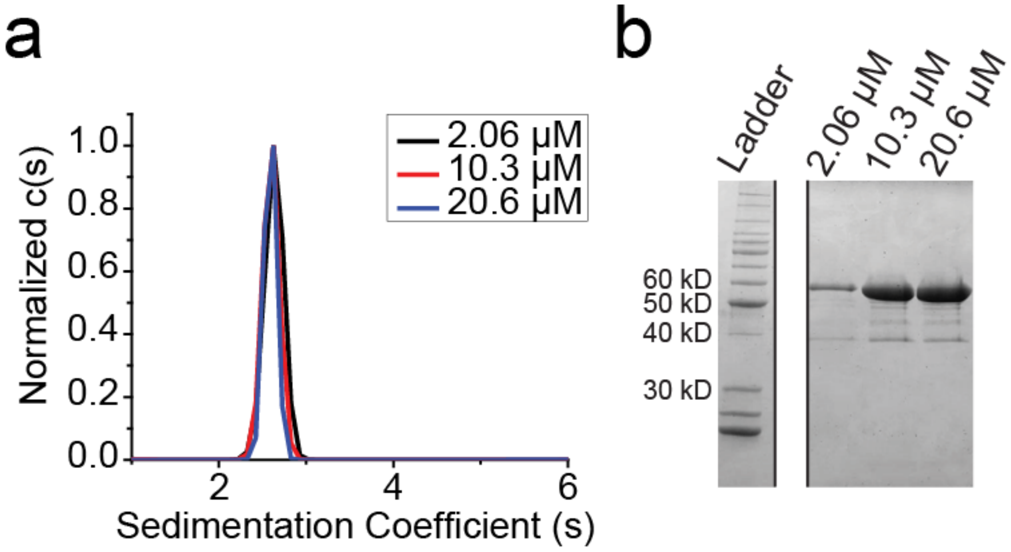


**Supplementary Figure 6. Bacterially produced PCDH1 EC1-4 C375S K398E does not dimerize in solution. (a)** AUC of PCDH1 EC1-4 C375S K398E shows only a monomeric peak for a range of concentrations between 2.06 μM and 20.6 μM. Normalized c(s) has been plotted against sedimentation coefficient (s). **(b)** SDS-PAGE analysis shows that samples for PCDH1 EC1-4 C375S K398E were pure and did not degrade after AUC. Samples were not loaded in equal amount for the SDS-PAGE.

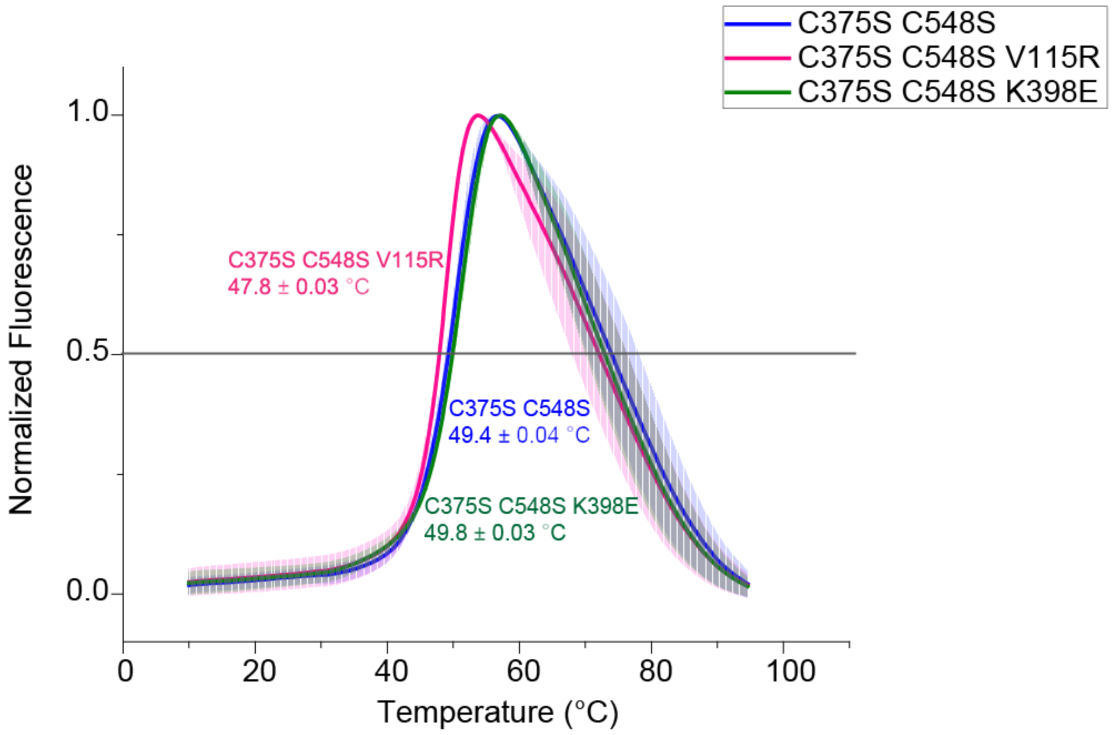


**Supplementary Figure 7. Melting temperatures of PCDH1 EC1-5 C375S C548S, EC1-5 C375S C548S V115R, and EC1-5 C375S C548S K398E determined using differential scanning fluorimetry show that their thermal stability is comparable.** The curves represent the average for each construct with vertical bars representing standard deviation from the mean ($n$ = 3).

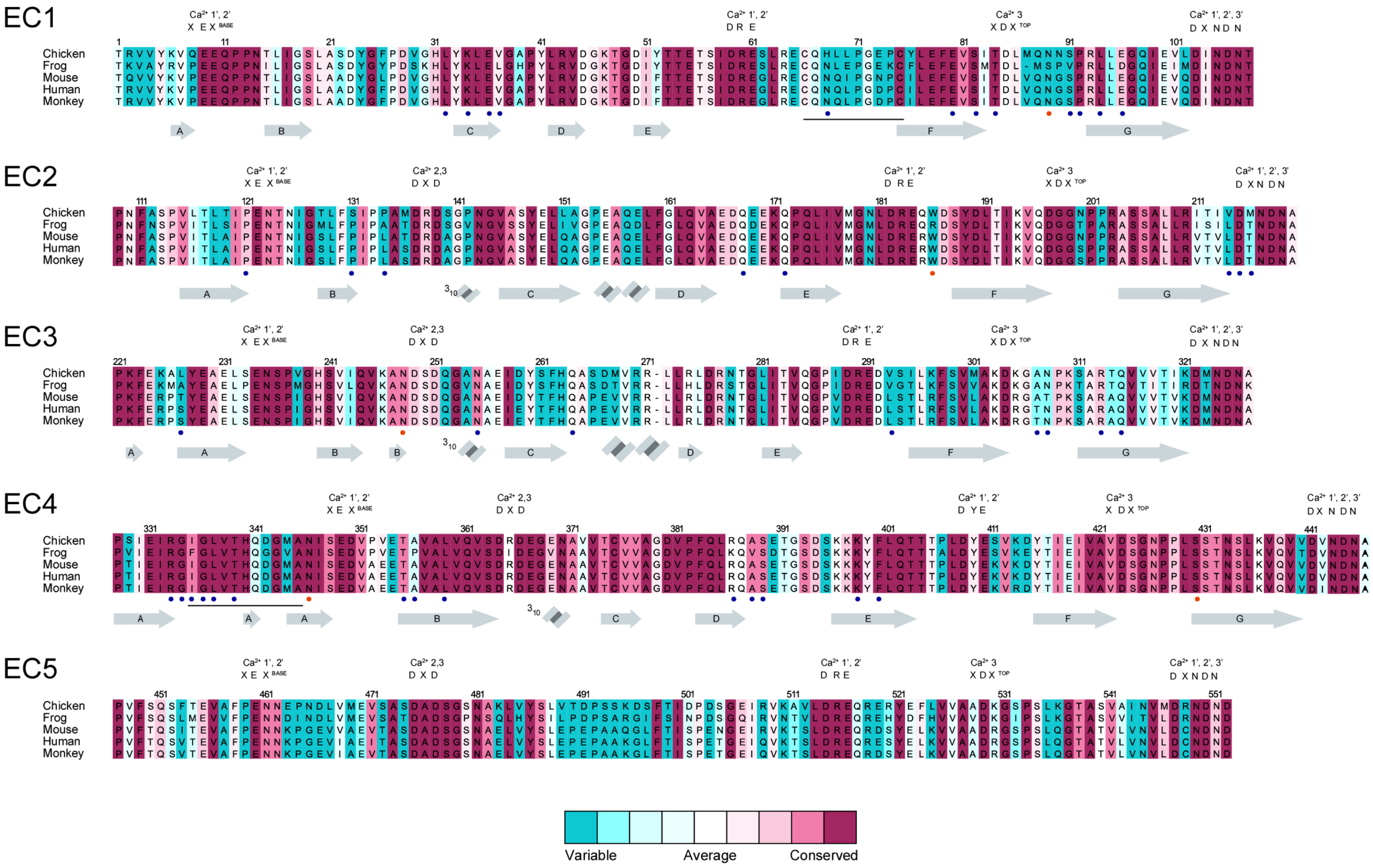


**Supplementary Figure 8. Sequence alignment of PCDH1 EC1-5.** Alignment of chicken, frog, mouse, human, and monkey sequences for PCDH1 EC1-5. Residues are colored according to conservation based on ConSurf and an alignment of sequences from 87 species. Conserved calcium-binding motifs are shown on top of the alignment and labeled. Interfacing residues are marked as blue dots. Predicted and observed glycosylation sites are marked as orange dots. The cysteine disulfide loop in EC1 and the δ insertion in EC4 are underlined. The accession numbers for the protein sequences of each species used are given in Supplementary Table 1.

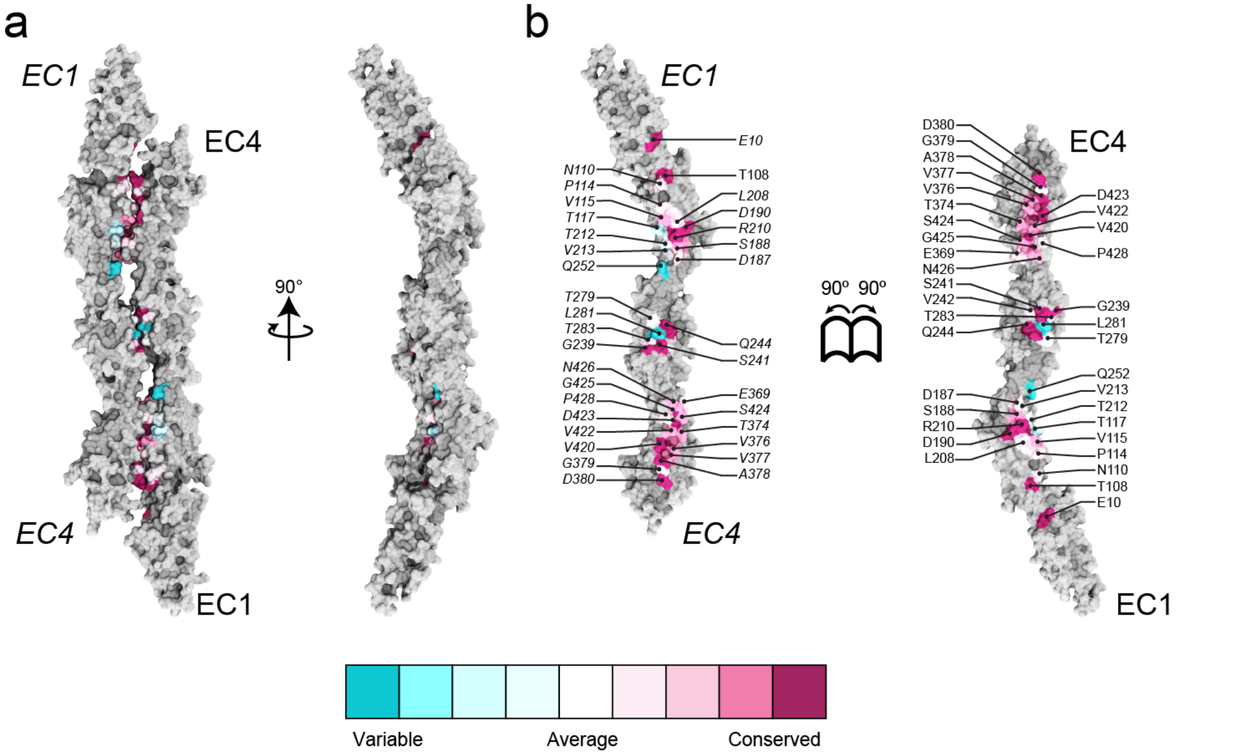


**Supplementary Figure 9. Conservation of residues at the crystallographic PCDH1-I2 interface. (a)** Molecular surface representation of PCDH1-I2 antiparallel dimer. Two perpendicular views are shown. **(b)** Interaction surface exposed with interfacing residues listed and colored according to sequence conservation among 87 species. The accession numbers for the protein sequences of each species used are given in Supplementary Table 1.

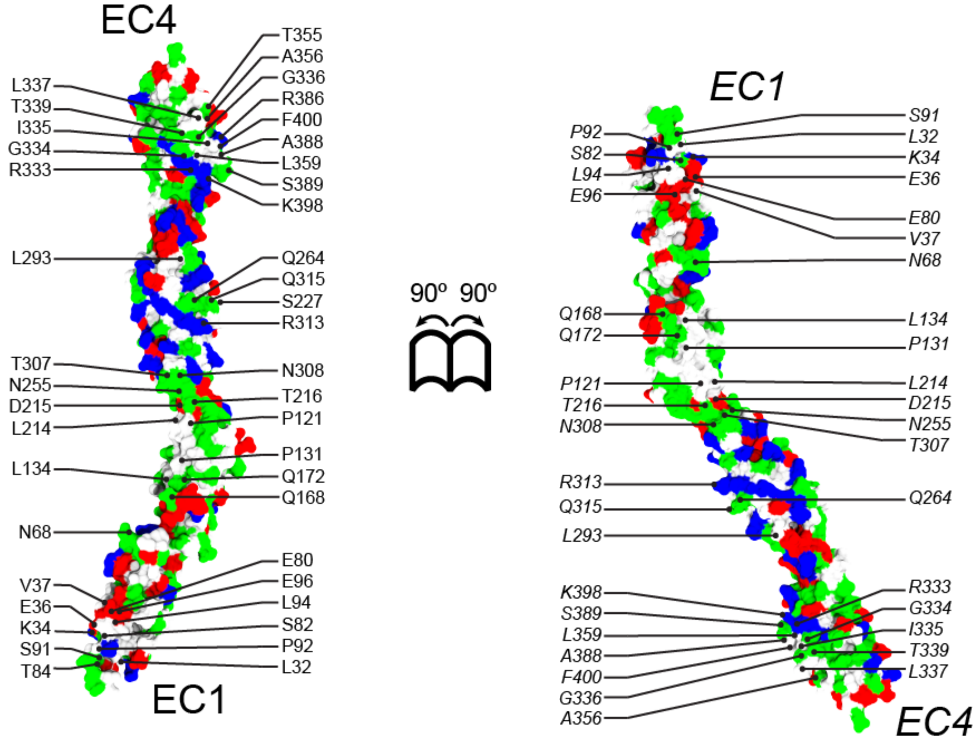


**Supplementary Figure 10**. **PCDH1 antiparallel EC1-4 dimer interface involves charged, hydrophilic, and hydrophobic residues.** Molecular surface representation of the PCDH1 EC1-4 antiparallel dimer with interfacial residues exposed and labeled. Surface is colored according to residue type (apolar: white; polar: green; negatively charged: red; positively charged and histidines: blue).

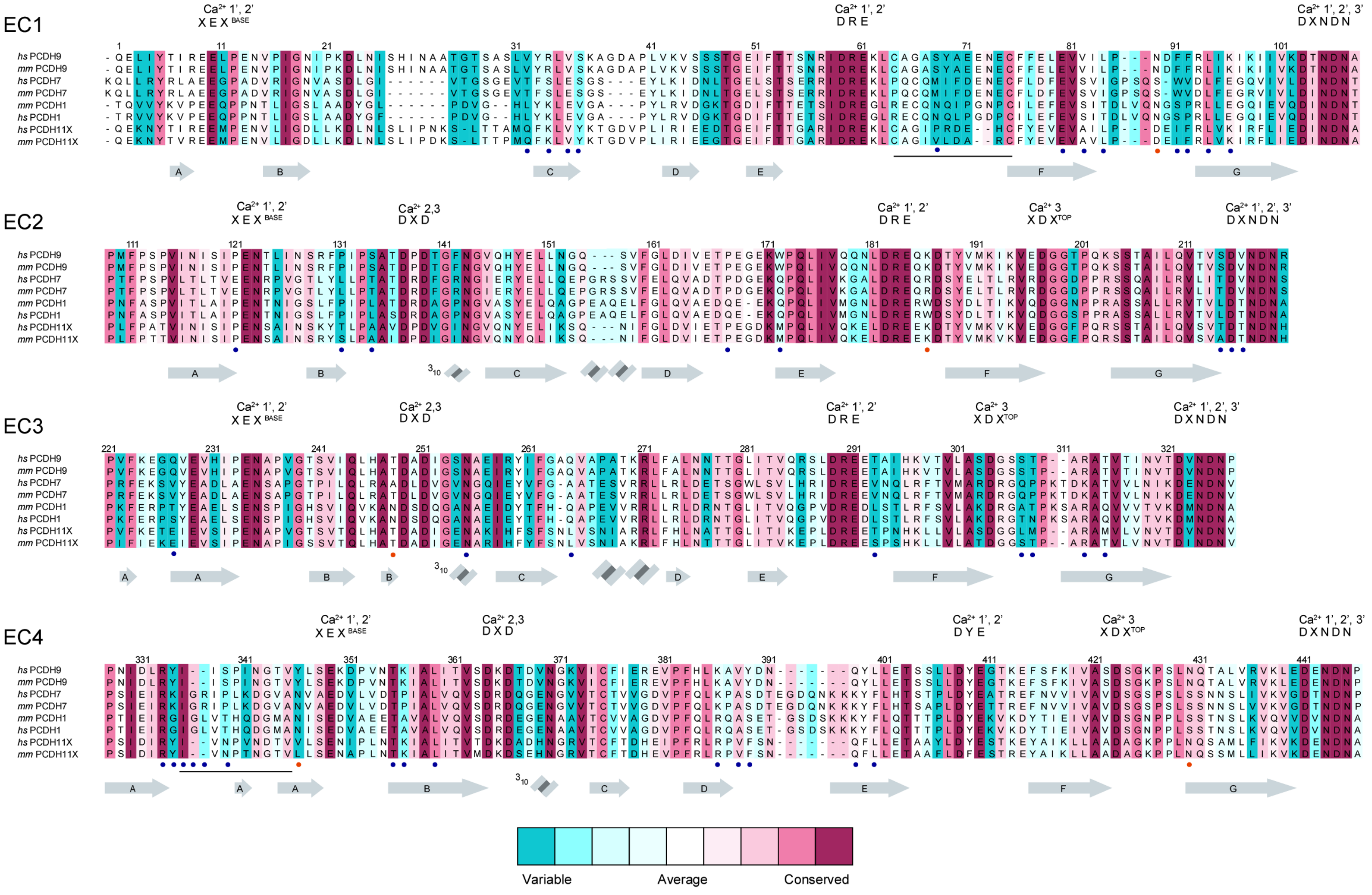

**Supplementary Figure 11. Sequence alignment of *hs* and *mm* PCDH9, PCDH7, PCDH1, and PCDH11X EC1 to EC4.** Calcium-binding motifs are labeled. Interfacing residues are marked as blue dots. Predicted and observed glycosylation sites are marked as orange dots. The cysteine disulfide loop in EC1 and the δ insertion in EC4 are underlined. The accession numbers for the protein sequences of each species used are given in Supplementary Table 2.

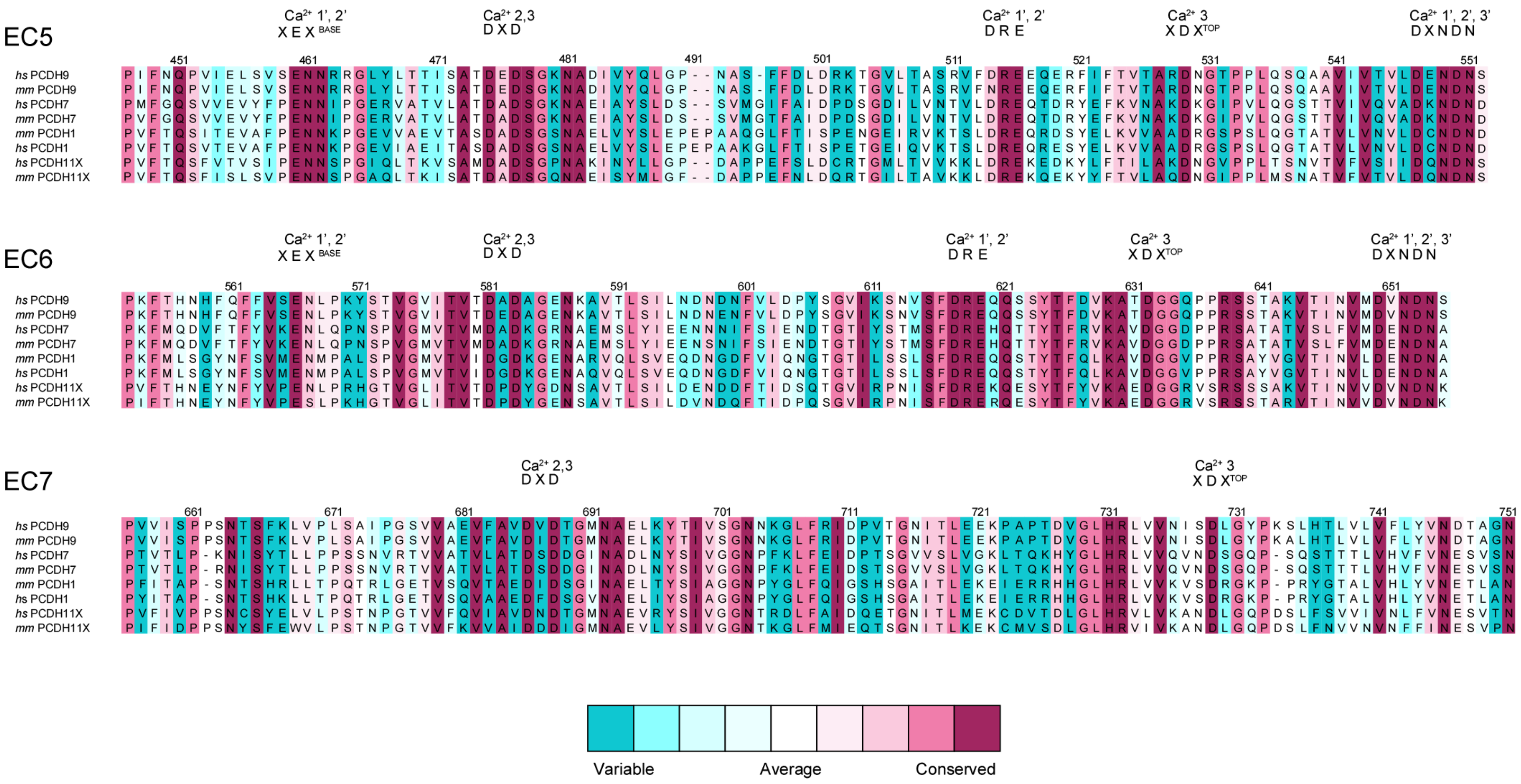


**Supplementary Figure 12. Sequence alignment of *hs* and *mm* PCDH9, PCDH7, PCDH1, and PCDH11X EC5 to EC7.** Calcium-binding motifs are labeled. The accession numbers for the protein sequences of each species used are given in Supplementary Table 2.

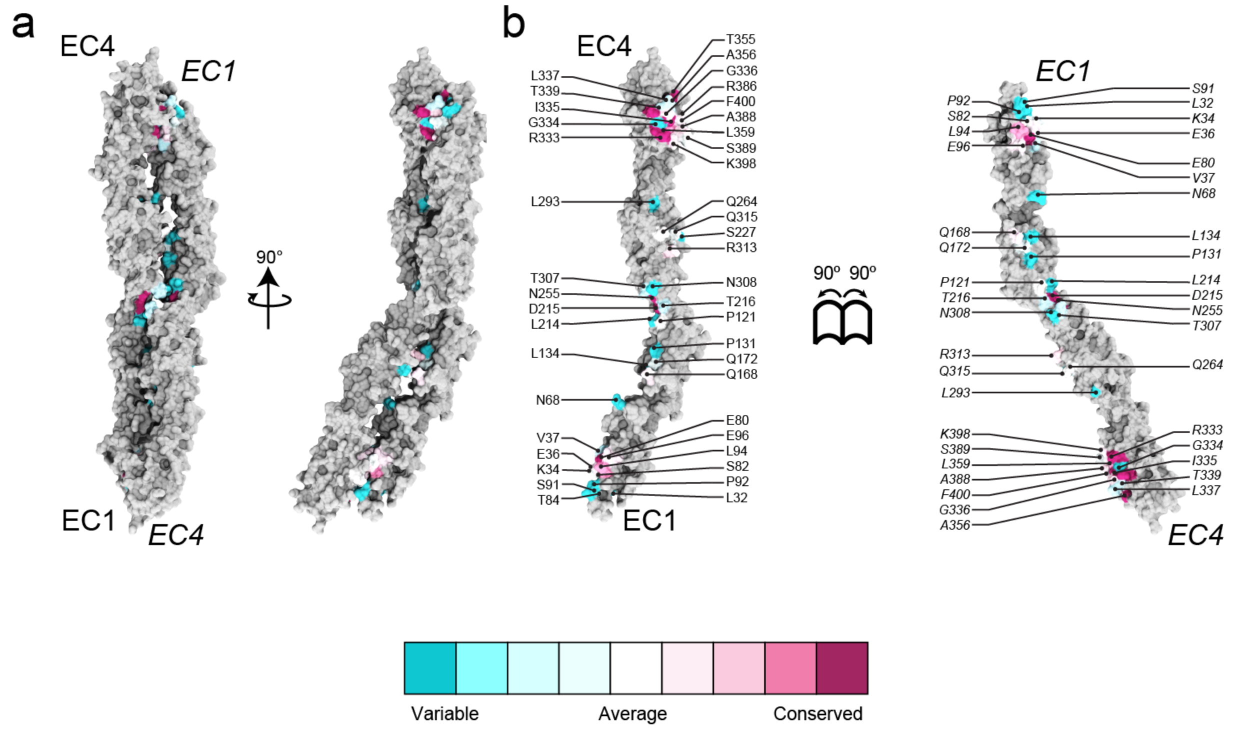


**Supplementary Figure 13. Conservation of residues among all δ1 protocadherins at the crystallographic PCDH1-I1 interface. (a)** Molecular surface representation of PCDH1-I1 antiparallel dimer. Two perpendicular views are shown. **(b)** Interaction surface exposed with interfacing residues listed and colored according to sequence conservation across different δ1 protocadherins. The accession numbers for the protein sequences of each species used are given in Supplementary Table 2.

δ1 Protocadherins Identity

| | | | | |
|---|---|---|---|---|
| ***hs*PCDH1** | 100% | | | |
| ***hs*PCDH7** | 53.54% | 100% | | |
| ***hs*PCDH9** | 45.27% | 43.92% | 100% | |
| ***hs*PCDH11X** | 42.91% | 38.13% | 58.98% | 100% |
| | ***hs*PCDH1** | ***hs*PCDH7** | ***hs*PCDH9** | ***hs*PCDH11X** |

δ1 Protocadherins Similarity

| | | | | |
|---|---|---|---|---|
| ***hs*PCDH1** | 100% | | | |
| ***hs*PCDH7** | 65.87% | 100% | | |
| ***hs*PCDH9** | 55.90% | 56.70% | 100% | |
| ***hs*PCDH11X** | 53.67% | 50.17% | 72.91% | 100% |
| | ***hs*PCDH1** | ***hs*PCDH7** | ***hs*PCDH9** | ***hs*PCDH11X** |

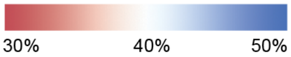


**Supplementary Figure 14. δ1 protocadherins are similar in sequence.** Identity and similarity matrices of δ1 protocadherins were calculated with the SIAS server using their EC1-7 extracellular domains.

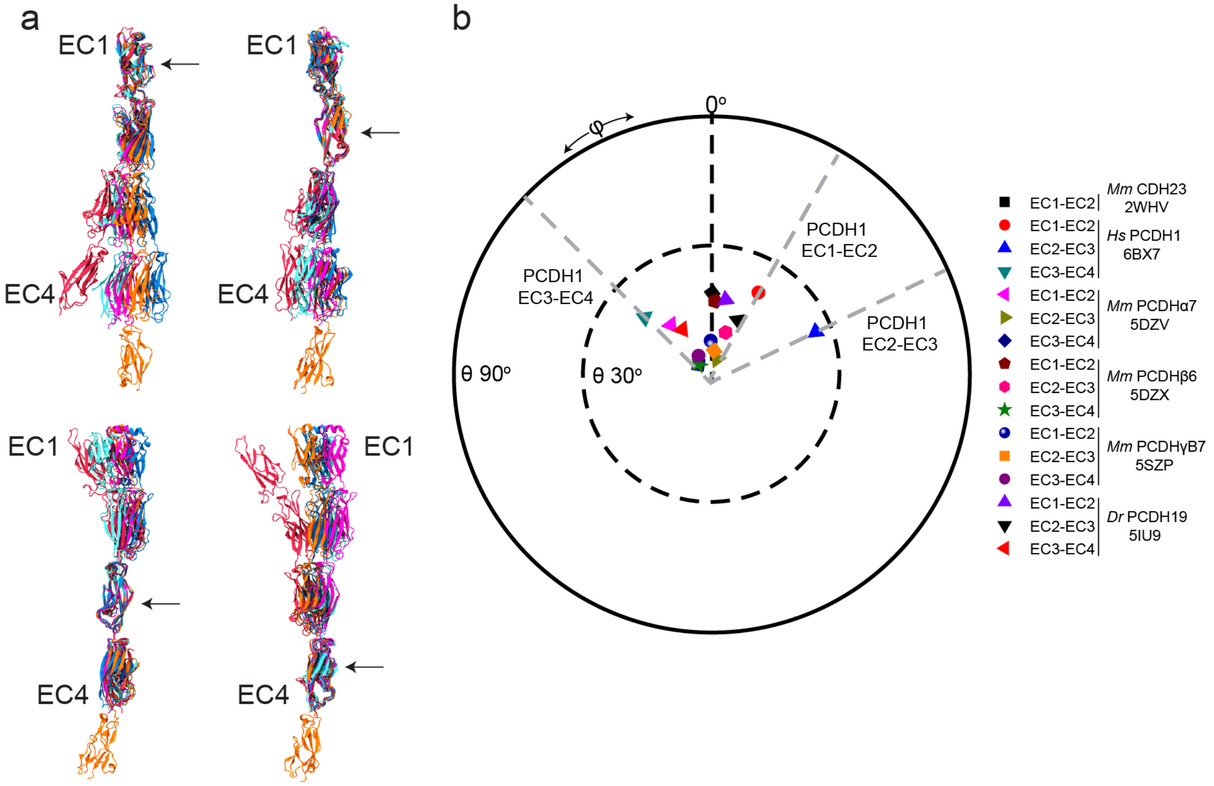


**Supplementary Figure 15**. **PCDH1 (δ1) is more curved than the clustered (α, β, γ) and PCDH19 (δ2) protocadherins. (a)** Alignments of ECs of PCDH1 (red), PCDHα7 (orange), PCDHβ6 (magenta), PCDHγB7 (blue) and PCDH19 (cyan) show that PCDH1 is more twisted than the others. The top left and right panels have EC1s and EC2s aligned, respectively. The bottom left and right panels have EC3s and EC4s aligned, respectively. Arrows indicate aligned EC. **(b)** Orientation of tandem EC repeats for listed cadherins shows that PCDH1 angles are different. The N-terminal EC repeat for labeled structures was used as reference and aligned to the *z*-axis. CDH23 EC1-2 was used to define φ = 0°. The azimuthal angle (φ) observed for PCDH1 EC2-3 is unique.

Supplementary Table 1. Accession numbers of the PCDH1 EC1-7 sequences of 87 species used for studying conservation of residues.

| S/N | Common name | Scientific name | Accession number |
|---|---|---|---|
| 1. | Human | *Homo sapiens* | NP_115796.2 |
| 2. | Mouse | *Mus musculus* | NP_083633.2 |
| 3. | Chicken | *Gallus gallus* | XP_015149085.1 |
| 4. | Rat | *Rattus norvegicus* | XP_006222627.1 |
| 5. | Cattle | *Bos taurus* | NP_001077124.1 |
| 6. | Orangutan | *Pongo abelii* | XP_024103364.1 |
| 7. | Northern white-cheeked gibbon | *Nomascus leucogenys* | XP_012360025.1 |
| 8. | Frog | *Xenopus tropicalis* | NP_001096175.1 |
| 9. | Dog | *Canis lupus familiaris* | XP_013963480.1 |
| 10. | Chimpanzee | *Pan troglodytes* | XP_009448113.1 |
| 11. | Horse | *Eqqus caballus* | XP_023473277.1 |
| 12. | Domestic ferret | *Mustela putoriusfuro* | XP_004744802.1 |
| 13. | Brandt's bat | *Myotis brandtii* | XP_014384591.1 |
| 14. | Golden hamster | *Mesocricetus auratus* | XP_012968155.1 |
| 15. | Pig | *Sus crofa* | XP_020940599.1 |
| 16. | Rhesus monkey | *Macaca mulatta* | XP_014996578.1 |
| 17. | Upper Galilee mountains blind mole rat | *Nannispalax galili* | XP_008833733.1 |
| 18. | Beluga whale | *Delphinapterus leucas* | XP_022445669.1 |
| 19. | Hawaiian monk seal | *Neomonachus schauinslandi* | XP_021557659.1 |
| 20. | Mongolian gerbil | *Meriones unguiculatus* | XP_021506942.1 |
| 21. | Helmeted guineafowl | *Numida meleagris* | XP_021266215.1 |
| 22. | Shrew mouse | *Mus pahari* | XP_021069920.1 |
| 23. | Koala | *Phascolarctos cinereus* | XP_020860723.1 |
| 24. | Great blue-spotted mudskipper | *Boleophthalmus pectinirostris* | XP_020785977.1 |
| 25. | Central bearded dragon | *Pogona vitticeps* | XP_020654882.1 |
| 26. | Whale shark | *Rhincodon typus* | XP_020375084.1 |
| 27. | Zebu cattle | *Bos indicus* | XP_019820689.1 |
| 28. | Chinese rufous horseshoe bat | *Rhinolophussinicus* | XP_019594645.1 |
| 29. | Great roundleaf bat | *Hipposiderosarmiger* | XP_019518573.1 |
| 30. | Australian saltwater crocodile | *Crocodylus porosus* | XP_019390369.1 |
| 31. | Gharial | *Gavialis gangeticus* | XP_019362111.1 |
| 32. | Leopard | *Panthera pardus* | XP_019280068.1 |
| 33. | Asian bonytongue | *Scleropages formosus* | XP_018592904.1 |
| 34. | Blue-crowned manakin | *Lepidothrix coronata* | XP_017685910.1 |
| 35. | White-faced capuchin | *Cebus capucinus imitator* | XP_017387745.1 |
| 36. | Natal long-fingered bat | *Miniopterus natalensis* | XP_016061056.1 |
| 37. | Egyptian rousette | *Rousettus aegyptiacus* | XP_015979646.1 |
| 38. | Japanese quail | *Coturnix japonica* | XP_015731461.1 |
| 39. | Brown-spotted pit viper | *Protobothrops mucrosquamatus* | XP_015672907.1 |
| 40. | Great tits | *Parus major* | XP_015496986.1 |
| 41. | Amur tiger | *Panthera tigris altaica* | XP_015391107.1 |
| 42. | Alpine marmot | *Marmota marmota marmota* | XP_015354329.1 |
| 43. | Gecko | *Gekko japonicus* | XP_015269557.1 |
| 44. | Cheetah | *Acinonyx jubatus* | XP_014918959.1 |

| | | | |
|---|---|---|---|
| 45. | Ruff | *Calidris pugnax* | XP_014811282.1 |
| 46. | Common starling | *Sturnus vulgaris* | XP_014745102.1 |
| 47. | Donkey | *Equus asinus* | XP_014705742.1 |
| 48. | Garter snake | *Thamnophis sirtalis* | XP_013908725.1 |
| 49. | North island brown kiwi | *Apteryxaustralis mantelli* | XP_013802378.1 |
| 50. | Domestic goose | *Anser cygnoides domesticus* | XP_013033983.1 |
| 51. | Ord’s kangaroo rat | *Dipodomys ordii* | XP_012875722.1 |
| 52. | Grey mouse lemur | *Microcebus murinus* | XP_012604524.1 |
| 53. | Coquerel’s sifaka | *Propithecus coquereli* | XP_012501284.1 |
| 54. | Ma’s night monkey | *Aotus nancymaae* | XP_012317787.1 |
| 55. | Sooty mangabey | *Cerocebus atys* | XP_011945991.1 |
| 56. | Pig-tailed macaque | *Macacanemestrina* | XP_011714527.1 |
| 57. | Golden eagle | *Aquila chrysaetoscanadensis* | XP_011569959.1 |
| 58. | Large flying fox | *Pteropus vampyrus* | XP_023390699.1 |
| 59. | Arabian camel | *Camelus dromedarius* | XP_010980451.1 |
| 60. | Bactrian camel | *Camelus bactrianus* | XP_010948845.1 |
| 61. | Bison | *Bison bison bison* | XP_010848704.1 |
| 62. | Damara mole-rat | *Fukomys damarensis* | XP_010612182.1 |
| 63. | Bald eagle | *Haliaeetus leucocephalus* | XP_010577476.1 |
| 64. | Hooded crow | *Corvus cornix cornix* | XP_010408469.1 |
| 65. | Golden snub-nosed monkey | *Rhinopithecus roxellana* | XP_010386558.1 |
| 66. | East African grey-crowned crane | *Balearicaregulorum gibbericeps* | XP_010299558.1 |
| 67. | White-throated tinamou | *Tinamusguttatus* | XP_010224802.1 |
| 68. | Chuck-will’s-widow | *Antrostomus carolinensis* | XP_010162738.1 |
| 69. | Macqueen’s bustard | *Chlamydotis macqueenii* | XP_010124265.1 |
| 70. | Chimney swift | *Chaetura pelagica* | XP_009999706.1 |
| 71. | Cuckoo roller | *Leptosomus discolor* | XP_009954693.1 |
| 72. | Reptile bird | *Opisthocomus hoazin* | XP_009942673.1 |
| 73. | Killdeer | *Charadrius vociferus* | XP_009879047.1 |
| 74. | South African ostrich | *Struthio camelus australis* | XP_009688499.1 |
| 75. | Dalmatian pelican | *Pelecanus crispus* | XP_009487536.1 |
| 76. | Crested ibis | *Nipponia nippon* | XP_009467776.1 |
| 77. | Adelie penguin | *Pygoscelis adeliae* | XP_009328306.1 |
| 78. | Emperor penguin | *Aptenodytes forsteri* | XP_009277624.2 |
| 79. | Common canary | *Serinus canaria* | XP_018770832.1 |
| 80. | Carmine bee-eater | *Merops nubicus* | XP_008938515.1 |
| 81. | Polar beer | *Ursus maritimus* | XP_008689915.1 |
| 82. | Przewalski’s horse | *Equus przewalskii* | XP_008517941.1 |
| 83. | Anna’s hummingbird | *Calypte anna* | XP_008501640.1 |
| 84. | Green monkey | *Chlorocebus sabaeus* | XP_008012957.1 |
| 85. | Elephant shark | *Callorhinchus milii* | XP_007883233.1 |
| 86. | Western European hedgehog | *Erinaceus europaeus* | XP_016041855.1 |
| 87. | Yangtze river dolphin | *Lipotes vexillifer* | XP_007468003.1 |

Supplementary Table 2. Accession numbers of δ1 protocadherin protein sequences used for studying conservation of residues across different family members.

| S/N | Name | Accession Number |
|---|---|---|
| 1. | *hs* PCDH1 | NP_115796.2 |
| 2. | *mm* PCDH1 | NP_083633.2 |
| 3. | *hs* PCDH7 | NP_001166994.1 |
| 4. | *mm* PCDH7 | NP_001116230.1 |
| 5. | *hs PCDH9* | NP_982354.1 |
| 6. | *mm* PCDH9 | NP_001074846.1 |
| 7. | *hs* PCDH11X | NP_116750.1 |
| 8. | *mm* PCDH11X | NP_001258738.1 |

Supplementary Table 3. Accession numbers of $\delta$1, $\delta$2, and clustered protocadherin protein sequences used for studying unique features in EC4 of PCDH1.

| S/N | Name | Accession Number |
|---|---|---|
| 1 | *hs* PCDH11Y | NP_116754.1 |
| 2 | *hs* PCDH8 | NP_002581.2 |
| 3 | *mm* PCDH8 | NP_067518.2 |
| 4 | *hs* PCDH10 | NP_116586.1 |
| 5 | *mm* PCDH10 | NP_001091640.1 |
| 6 | *hs* PCDH17 | NP_001035519.1 |
| 7 | *mm* PCDH17 | NP_001013775.2 |
| 8 | *hs* PCDH18 | NP_061908.1 |
| 9 | *mm* PCDH18 | NP_569715.3 |
| 10 | *hs* PCDH19 | NP_001171809.1 |
| 11 | *mm* PCDH19 | NP_001098715.1 |
| 12 | *hs* PCDH$\alpha$7 | NP_061733.1 |
| 13 | *mm* PCDH$\alpha$7 | NP_034087.1 |
| 14 | *hs* PCDH$\alpha$4 | NP_061730.1 |
| 15 | *mm* PCDH$\alpha$4 | NP_031792.1 |
| 16 | *hs* PCDH$\beta$6 | NP_061762.2 |
| 17 | *mm* PCDH$\beta$6 | NP_444361.1 |
| 18 | *hs* PCDH$\beta$8 | NP_061993.3 |
| 19 | *mm* PCDH$\beta$8 | NP_444363.1 |
| 20 | *hs* PCDH$\gamma$B7 | NP_061750.1 |
| 21 | *mm* PCDH$\gamma$B7 | NP_291057.1 |
| 22 | *hs* PCDH$\gamma$B3 | NP_061747.2 |
